# Gut Microbiome Analysis In Adult Tropical Gars (*Atractosteus tropicus*)

**DOI:** 10.1101/557629

**Authors:** Roberto Méndez-Pérez, Rodrigo García-López, J. Santiago Bautista-López, Jorge F. Vázquez-Castellanos, Emyr S. Peña-Marín, Rafael Martínez-García, Verónica I. Domínguez-Rodríguez, Randy H. Adams-Schroeder, Eduardo Baltierra-Trejo, Carolina Melgar Valdés, Andrés Moya, Carlos A. Alvarez-González, Rodolfo Gómez-Cruz

## Abstract

Tropical gar (*Atractosteus tropicus*), is freshwater and estuarine fish that has inhabited the Earth since the Mesozoic era, undergoing limited physiological variation ever since. This omnivorous fish is endemic to southern Mexico and part of Central America. Besides its recognized cultural and scientific relevance, the species has seen remarkable growth in its economic impact due to pisciculture. Previous studies have highlighted the role of microbial communities in fish, particularly those in the gut microbiome, in maintaining their host homeostasis or disease. In this study, we present the first report of the whole taxonomic composition of microbial communities in gut contents of adults’ *A. tropicus*, by sex (female/male) and origin (wild/cultivated). Using culture-independent techniques, we extracted metagenomic DNA that was used for high throughput 16S rDNA profiling by amplifying the V4 – V5 hypervariable regions of the bacterial gene. A total of 364,735 total paired-end reads were obtained on an Illumina MiSeq sequencing platform, belonging to 508 identified genera, with the most and least abundant are *Cetobacterium, Edwardsiella, Serratia, Clostridium sensu stricto, Paludibacter* and *Campylobacter, Snodgrassella, Albirhodobacter, Lentilitoribacter*, respectively. We detected that, by sex and origin, Proteobacteria, Fusobacteria, Firmicutes and Bacteroidetes phyla are the core gut microbiome of the adults’ *A. tropicus*. We discover the Deinococcus-Thermus phylum sequence, wildtype males only, with extremophile capacity in another freshwater fish. We also identified the species *Lactococcus lactis* strains CAU929 and CAU6600, Cp6 and CAU9951, *Cetobacterium* strain H69, *Aeromonas hydrophila* strains P5 and WR-5-3-2, *Aeromonas sobria* strain CP DC28 and *Aeromonas hydrophila* with probiotic potential in aquaculture within the three dominant phyla, especially in wild-type organisms.

## 1. Introduction

Aquaculture produces 76.6 million tons of fish for human consumption and economically ensures the livelihood of 10 to 12 % of the world population (FAO 2014, 2017). It has become the fastest growing food sector in the world, with an average annual growth rate of 8.9 % since 1970 (Subasinghe 2005) and there is a growing global trend towards diversifying the spectrum of cultured aquatic species. Biological diversity in Latin America is one of the richest on the planet, including an important variety of its freshwater ichthyofauna (Flores-Nava y Brown 2010). Some of the greatest challenges for aquaculture during this millennium have been the creation of integral studies on pisciculture-revellant endemic species and the development of technologies that may allow for controlled production of these fish in a profitable, innocuous and environmentally-conscious approach (Márquez-Couturier y Vázquez-Navarrete 2015). Studies in other fish have explored the bacterial populations in varying habitats such as the skin, gills, eggs and gut emicrobiome (GMB) and the way they finfluence the host’s general health and physiology (MacFarlane *et al*. 1986, Cahill 1990, Ringø & Tabachek 1995, Givens 2012, Austin & Austin 2016). They have reported a large variation of microbiota among the different niches and between species with the GMB as one of the most studied due to its high microorganism concentration. Of interest for quaculture aquaculture-relevant species are outbreaks of viral, bacterial and fungal infections as they may cause devastating economic losses worldwide due to poor environmental conditions in the farms, unbalanced feeding, generation of toxins and genetic factors (Martínez-Cruz *et al.* 2012).

Tropical gar (*Atractosteus tropicus*, also known as pejelagarto) is freshwater and brackish fish species of the Lepisosteidae family, which has an ample fossil record since the Cretaceous period of the Mezosoic Era (Wiley 1976, Reséndez-Medina & Salvadores-Baledón 1983). The morphology of these species has remained mostly unaltered, with current specimens having lengths between 1.0 and 1.2 m and weighing between 1000 and 3000 g in the wild. In their natural habitat, this species exists in coastal wetlands of the tropical rainy areas of southeastern Mexico, Belize, Guatemala, El Salvador, Honduras, Nicaragua and Costa Rica (Willey 1976, Bussing 1998, Miller *et al*. 2005, Nelson 2006). It is an omnivore, feeding on other fish, decomposing organic matter, crustaceans, plants, etc., depending on the availability, preferring carnivorous habits. This specially secluded species has seen a drastic decrease in wild populations, caused by anthropogenic activities that have led to the loss of habitats and severe ecological alterations (Méndez-Marín *et al*. 2012). Currently, *A. tropicus* is cultivated in fish farms for human consumption in Mexico (Márquez *et al*. 2015).

The gastrointestinal tract of A. tropicus, is formed by the buccopharyngeal, esophagus, stomach, gut, pyloric blind, rectum and anus, which is rapidly developing during the larval period (Frías-Quintana *et al*. 2015). Having absorbed and secretory function, its intestine is a long tube, narrower than the stomach with an inner epithelial mucosa that forms long, and numerous folds comprised of high columnar cells with clearly defined brush border (Márquez-Couturier *et al*. 2006). Having absorbed and secretory function, its intestine is a long tube, narrower than the stomach with an inner epithelial mucosa that forms long, and numerous folds comprised of high columnar cells with clearly defined brush border (Márquez-Couturier *et al*. 2006). In the anterior and middle regions, long folds divide these regions into a series of compartments called a spiral valve. The shorter posterior region forms the rectum, which opens in the anus. The external muscle in the intestine is thinner than in the stomach. Early juveniles 20 days after hatching show a GMB like that of adults. toBefore colonization, microorganisms may access the GMB through food and water intake during the larval phase (Núñez de la Rosa 2011). The high abundance of certain groups of gastrointestinal bacteria in fish, when compared to the microbial composition of the surrounding water suggests that GMB poses as a unique niche for a specific, but diverse group of bacteria (Cahill 1990, Givens 2012). It has been reported that GMB fish has between 10^7^ and 10^11^ bacteria per gram of feces (Nayak 2010). GMB fish plays an important role and directly influences the host’s nutrition and general homeostasis. Normal GMB fish may contain both beneficial and potentially pathogenic bacteria. The loss of the microbiota equilibrium (dysbiosis) has been reported to impact the host’s physiological state, potentially compromising immunity, growth, general development as well as the overall quality of the aquaculture production due to an increase in fish morbidity and mortality (Núñez de la Rosa 2011, Al-Harbi and Udding, 2005).

Prior to 2005, microbial studies on fish relied exclusively on culture techniques for enumerating and identifying bacteria (Newman *et al*. 1972, MacFarlane *et al*. 1986, Spanggaard *et al*. 2000, Aschfalk & Müller 2002, Verner-Jeffreys *et al*. 2003, Al-Harbi & Uddin 2004, Martin-Antonio *et al*. 2007, Skrodenyté 2007), providing valuable information as isolates provide a suitable framework for studying individual culturable strains on a finer level but limited appreciation of the whole spectrum in microbial communities. In 2007, Izvekova and collaborators reviewed GMB fish studies published between 1929 and 2006 reporting 73 different bacteria. This is because culturable bacteria typically account for less than 1 % of the cells that are present by direct microscopic enumeration (Ferguson *et al*. 1984, Head *et al*. 1998).

The advent of culture-independent metagenomic studies during the mid-2000s enabled the simultaneous analysis of complex genomic information contained in a hundreds of microbial species in a single niche (Nielsen *et al*. 2014). These techniques circumvent most culturing requirements of microorganisms, avoiding collection and sampling biases, effectively representing the actual diversity of a microbial community. Under intensive production conditions for sustainable aquaculture, aquatic species are subjected to high-stress conditions, leading to an increased incidence of diseases that decrease productivity (Bondad *et al*. 2005). It has been proposed that microbiota dysbiosis may be avoided through the regulation of their microbiota (Verschuere *et al*. 2000).

Microbial studies in aquaculture focus on the understanding of the symbiotic or antagonistic interactions between microbes and their eukaryotic hosts such as fish, crustaceans and molluscs. In this sense, metagenomics can provide a deeper understanding of these relationships through information revealed by sequencing microbial DNA extracted from specific niches within host organisms and, in the case of 16S profiling, taxa are representative of the medium (Suttle 2007, Gianoulis *et al*. 2009). This latter consists of surveying the 16S rRNA gene of all present microorganisms as this marker is found in all prokaryotes with enough mutations to discern each taxon. Formerly, some bacteria had been difficult to isolate because some of them are obligate intracellular microorganisms that could only be cultured in semi-aqueous and/or cellular culture media (Avila-Villa *et al*. 2011). Current sequencing platforms and bioinformatic tools enable the research on the diversity of intracellular bacteria, but also to ignore culture other culture requirements altogether as the DNA from the whole community are sequenced as is, with majority and accessory species, in order to elucidate the relevant community configurations for the improvement of aquaculture techniques. Consequently, the objective of this research was to explore the bacterial composition in the gut of adults’ *A. tropicus*, by analyzing the 16S rRNA gene profiles from adult male and female organisms, cultivated and wild, for biotechnological-relevant applications future.

## 2. Materials and methods

### Specimen collection

Fifteen totals live adults’ *A. tropicus* were collected for the study, 7 of them were cultivated in the Tropical Aquaculture Laboratory, <u>Research Center for Conservation and Sustainable Use of Tropical Resources</u> (CICART) at <u>Biological Sciences Academic Division</u> (DACBiol), <u>Juárez Autonomous University of Tabasco State</u> (UJAT), Mexico, and 8 wildtype, with an average weight and length of 5 kg and 1 m, respectively. This specimens were provided by fishermen of the municipalities of Nacajuca (<u>18°14′50″N 92°49′58″O</u>; n female, m male) and Centla (<u>18°20′00″N 92°30′00″O</u>; n female, m male), Tabasco, Mexico (Figure 1). All specimens were cut lengthwise under sterile conditions to remove the intestine for extracting its contents with scissors disinfected in absolute ethyl alcohol, for storage at −20 °C in a 2 mL Eppendorf tubes.

**Figure 1.**
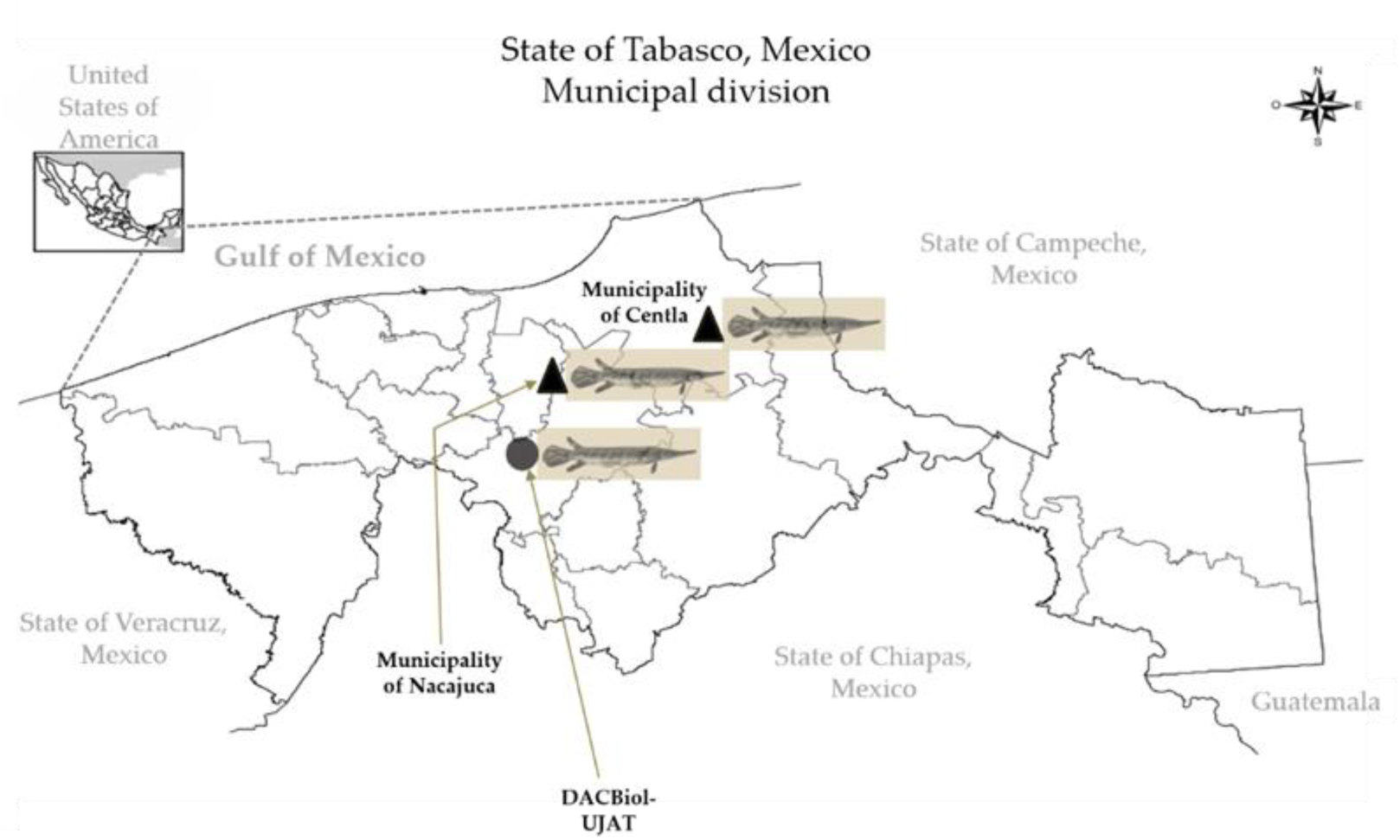
Collection sites of wild (triangle figures) and cultivated (circle figure) of the adults’ tropical gars (*Atractosteus tropicus*)

### Samples collection

All organisms were sacrificed according to the protocol was approved by Secretaría de Agricultura, Ganadería, Desarrollo Rural, Pesca y Alimentación (NOM-062-ZOO-1999) on 18 June 2001 by percussion stunning method, consisting of striking a quick blow on the head of the fish manually with a bat, to obtain fresh samples, squeezing the GMB and obtaining the feces separately, considering the geographical origin and sex by sterile conditions.

### DNA extraction, pyrosequencing and sequence for metagenomic analysis

200 mg of fresh feces were used and 2 mL were enough to ensure the target DNA with a capacity of the Eppendorf tubes selected. A whole genomic DNA (gDNA) extraction was carried out for each sample with a QIAamp DNA Stool Mini Kit (Qiagen, Valencia, CA, USA), following the manufacturer’s instructions. DNA integrity and concentration was evaluated by electrophoresis submerged with 1.2 % agarose gel and by spectrophotometry with a GenovaNano spectrophotometer (Jenway, Stone, Staffs, UK), respectively. Universal primers 533F (5’-GTGCCAGCAGCCGCGGTAA-3’) (Weisburg 1991) and 909R (5’-CCCCGYCAATTCMTTTRAGT-3’) (Tamaki *et al*. 2011) were selected for the amplification a region including the V4 and V5 hypervariable region in the16S rRNA gene. PCR amplifications were carried out using Phusion High-Fidelity DNA Polymerase (Finnzymes OY, Espoo, Finland; Klindworth *et al*. 2013) and conditions were as follows: Library preparation and high throughput sequencing was performed through Research & Testing Labs (Lubbock, Texas, USA) services, with an Illumina MiSeq platform, using 16S rDNA profiling by amplifying the V4 – V5 hypervariable regions of the bacterial gene, reagents sequencing for 2x 300 bp paired-end reads.

### Bioinformatic analysis

The sequences were subjected to the standard quality protocols, which included the sequencing adapters removal with the Cutadapt tool (1.18), the too short and low-quality sequences filtering based on the Phred quality score implemented in the PRINSEQ tool.lite (0.20.4). The paired-end were concatenated with -fastq_join command of the USEARCH program (V.11) and the chimeras were removed with – uchime2_ref command of the same program, using as reference the Gold database (microbiomeutil-r20110519). The sequences were clustered in OTUs with 97 % identity by usearch61_ref algorithm, using the SILVA132 database as a reference through the pick_otus.py command in QIIME (1.9) and OTUs were filtered with 3 or fewer sequences.

Alpha diversity was measured through the Chao1, Shannon and P. D. (Phylogenetic Diversity) diversity indexes, which were calculated by alpha_rarefaction.py command in QIIME (1.9). The Alpha diversity among samples and origin of the samples (wild and farmed) were compared using Kruskal-Wallis statistic (Multiple range test, Bonferroni post-hoc) and Mann-Whitney W statistic (Wilcoxon), respectively, both implemented in the ggpubr library of R software (V.3.4.4).

Beta Diversity was analyzed using a PCoA constructed from the Unweighted UniFrac distance, calculated with the beta_diversity_through_plots.py command from QIIME (1.9), the difference among groups was evaluated by ANOSIM statistical test, and the main coordinates of the PCoA were graphed in the factoextra library in R (V.3.4.4) and Differential Analysis of Abundance was compared the abundance patterns of the taxa between the groups of samples through LEfSe (Linear discriminant analysis effect size).

### Phylogenetic analysis

A sequences base was constructed according to certain genera with probiotic potential in fish that we find in the literature sought (Table 1), 16S rRNA gene sequences were obtained from the NCBI GenBank database. The 6,266 sequences obtained were clustered to eliminate identical sequences and generate a table of OTUs with 97 % similarity, using the usearch61 algorithm of QIIME (1.9). The database was indexed (Query Sequence) to be used in a local blast with the BLAST + tool (NCBI). The local blast was performed using the sequences of the samples (Subject sequences) clustered at 97 % as mentioned in the section on the processing of the sequences. The identity percentage was adjusted to 99 % (-perc_identity) and a coverage of 70 % (-qcov_hsp_perc 70). For phylogenetic analysis, the blast resulting Query and Subject sequences were subjected to a multiple alignment with the ClustalW algorithm, in MEGAX (Molecular Evolutionary Genetics Analysis), and the phylogenetic relationships were inferred by the Neighbor-joining method in MEGAx, with the predefined settings.

## 3. Results

The pyrosequencing method of the Illumina MiSeq platform was evaluated that uses the Research & Testing Labs (Lubbock, Texas, USA) services, applying 16S rRNA in V4 – V5 hypervariable regions. A total of 364,735 sequencing reads were generated, organized by sex and origin, according to Table 2. We can visualize in Table 2 that readings obtained of wildtype male organisms are far greater than of wildtype and cultivated females, and cultivated males. Sequences were clustered into 16,503 different OTUs (97 % identity), which were classified into 11 phyla, 22 class, 37 order, 86 families and 179 genera. Table 2 shows that about twice as many reads correspond to wild male.

### Microbiota composition

The cultivated organisms, wildtype females and males’ samples are dominated by the *Fusobacteria* (37.32 %), *Proteobacteria* (30.49 %) and *Firmicutes* (18.55 %) phyla, respectively (Figure 2a). In wildtype females the most abundant phyla were Proteobacteria (0.70 ± 0.28 SD), Firmicutes (0.38 ± 0.32 SD) and Fusobacteria (0.24 ± 0.29 SD) and in cultivated females are Proteobacteria (0.64 ± 0.20 SD), Fusobacteria (0.54 ± 0.25 SD) and Bacteroidetes (0.45 ± 0.006 SD). In wildtype and cultived males, the most abundant phyla are Fusobacteria (0.63 ± 0.39 SD), Proteobacteria (0.40 ± 0.26 SD) and Fusobacteria (0.76 ± 0.36 SD), Proteobacteria (0.19 ± 0.21 SD), respectively. Actinobacteria (0.16 ± 0.17 SD) and Deinococcus-thermus (0.003 ± 0.006 SD) were identified only in wildtype females and wildtype males, respectively. The same tendency is observed in male and female specimens, since the dominant phyla in both wild type and cultivated ones are Fusobacteria (0.63 ± 0.39 SD), Proteobacteria (0.70 ± 0.28 SD) and Fusobacteria (0.76 ± 0.36 SD) and Proteobacteria (0.64 ± 0.20 SD), respectively (Figure 2a). At the genus level, the most abundant in wildtype females are *Edwardsiella* (0.32 ± 0.53 SD) and *Cetobacterium* (0.24 ± 0.29 SD), and in cultivated females are *Serratia* (0.57 ± 0.17 SD) and *Cetobacterium* (0.54 ± 0.25 SD). In wildtype males, the most abundant genera are *Cetobacterium* (0.63 ± 0.39 SD) and *Clostridium sensu stricto* (0.32 ± 0.49 SD) in cultivated males *Cetobacterium* (0.75 ± 0.29 SD) and *Edwardsiella* (0.22 ± 0.30 SD) (Figure 2b). In the case *Cetobacterium* (37.34 %) and *Edwardsiella* (10.55 %) genera are the most abundant in cultivated organisms and wildtype females’ samples, respectively (Figure 2b); whereas *Clostridium sensu stricto* (13.53 %) was more abundant in wildtype males’ samples. The cultivated and wildtype organisms’ samples are dominated by the Fusobacteria (38.95 %) and Firmicutes (19.57 %) phyla, respectively (Figure 2a). For the *Cetobacterium* (38.95 %) and *Clostridium sensu stricto* (15.09 %) genera they are more abundant in the cultured organisms and samples of wildtype organisms’ samples, respectively (Figure 2b).

**Figure 2.**
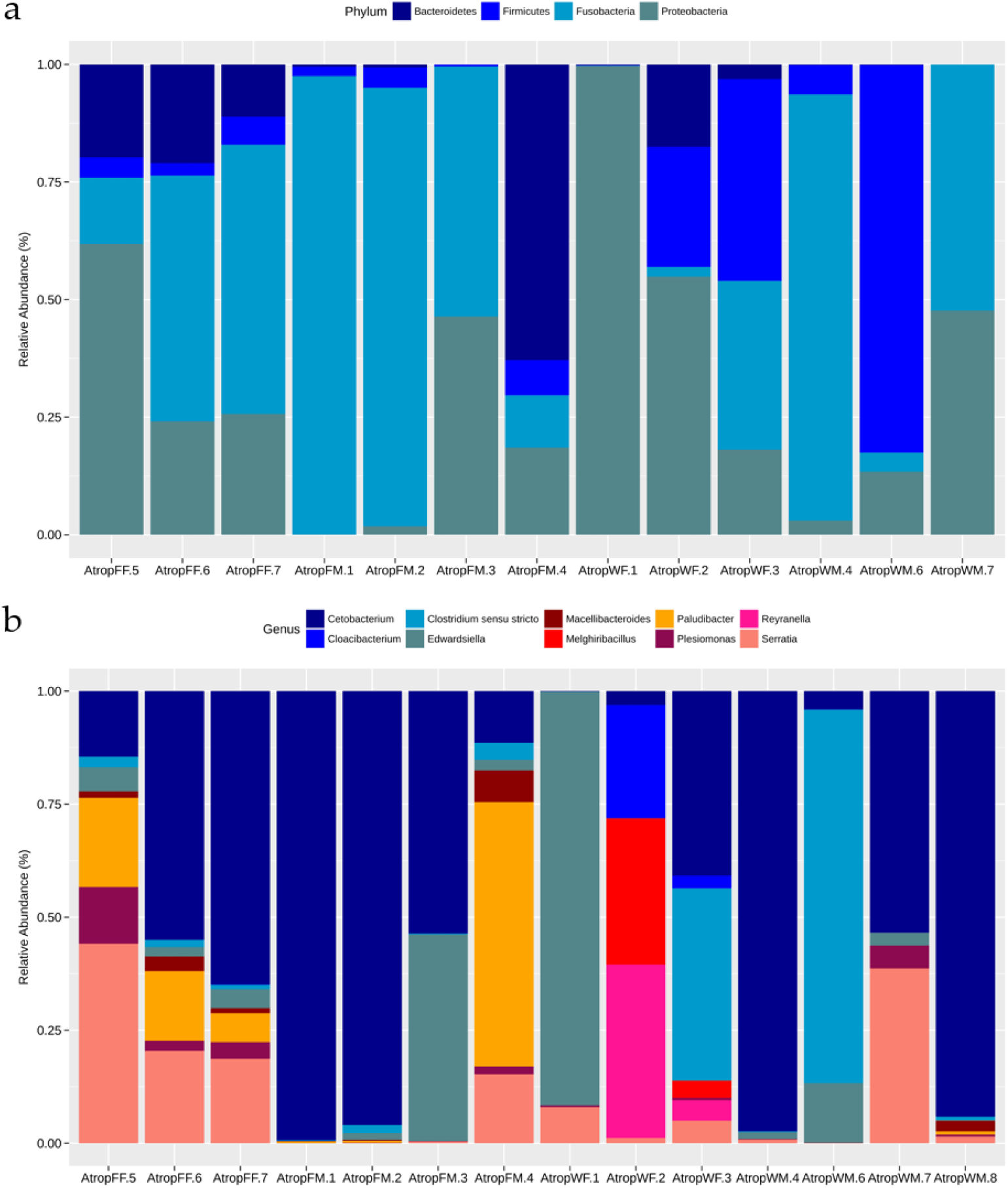
Taxonomic summary at different levels of relative bacterial abundance. (a) Phyla and (b) Genus. First letter W: wild or F: Farming individuals. Second letter sex M: male or F: female. Only taxa with relative abundance ≥ 1 % are shown.

Likewise, the core microbiome composition at phylum and genus taxonomic levels, on a per-sample basis of *adults’ A. tropicus* gut (figure 3), shows that the most abundant phyla per sample were Fusobacteria (47.01 %), Proteobacteria (29.02 %) and Firmicutes (12.90 %) (Figure 3a), while the most abundant genera were *Cetobacterium* (47.01 %), *Edwardsiella* (12.00 %) and *Serratia* (10.55 %) (Figure 3b). Although the primers are designed only to 16S rRNA gene amplify of bacteria, OTUs belonging to the archaea domain were also identified, represented by the genera *Salinirubrum, Salinigranum* and *Methanoculleus*, which together represent about 0.01 % of the total abundance.

**Figure 3.**
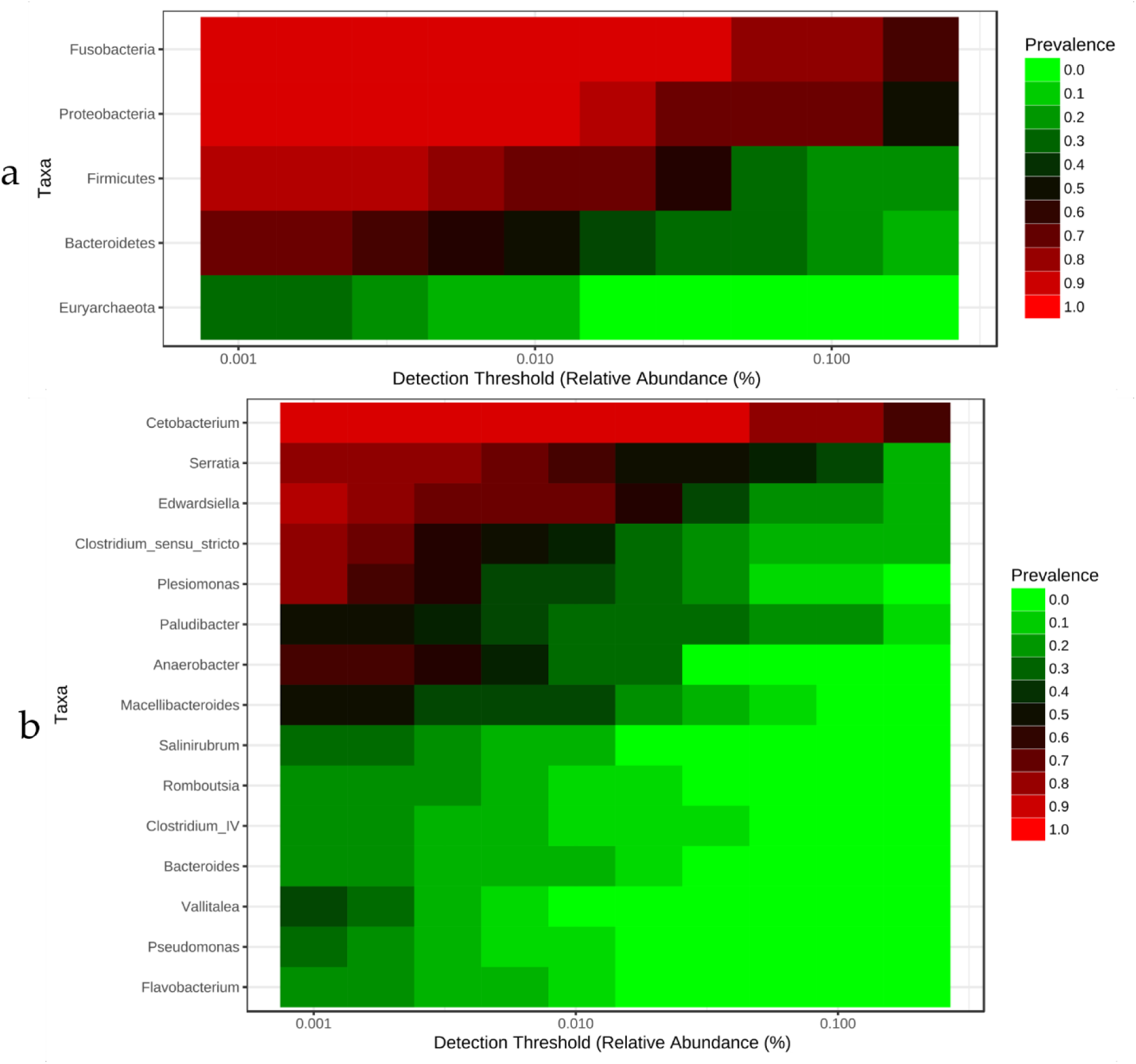
Core microbiome in gut of cultivated/wild and female/male adults’ *A. tropicus*, at taxonomic class-level of (a) phylum and (b) genus.

### Alpha diversity

In this work, diversity, dominance and richness were evaluated using the Shannon index about the total number of OTUs. Microbiota composition is influenced by source and sex factors. The Shannon index indicates that the microbial types in adults’ *A. tropicus* are more evenly distributed and thus may be more diverse in wild (5.32 ± 0.13 SD) and cultivated (3.75 ± 0.09 SD) female individuals than in the wild (3.14 ± 0.07 SD) and cultivated (3.38 ± 0.08 SD) male individuals (Table 3). We found significant differences in diversity indices and only some samples were not significant. The wild and cultivated females turned out to be the most diverse, but by origin, only the wild samples were the most diverse. In a total of 50 taxa, significant differences can be observed between cultivated and wild conditions.

### Beta diversity

Bacterial communities beta diversity associated to TGI of adults’ *A. tropicus* in origin conditions were measured through the ordination analysis from Principal Coordinates Analysis (PCoA), using Unweighted unifrac distances (UniFrac). The analysis produced an ordination of the dissimilarities, where similar individuals are close to one another and dissimilar ones are more distant (Figure 4). All ordination analysis showed a clear separation between wild and cultivated individuals samples. ANOSIM statistic (R: 0.60933; p-value <0.002) found significant statistical differences by origin, between the cultivated and wild groups (Figure 4).

**Figure 4.**
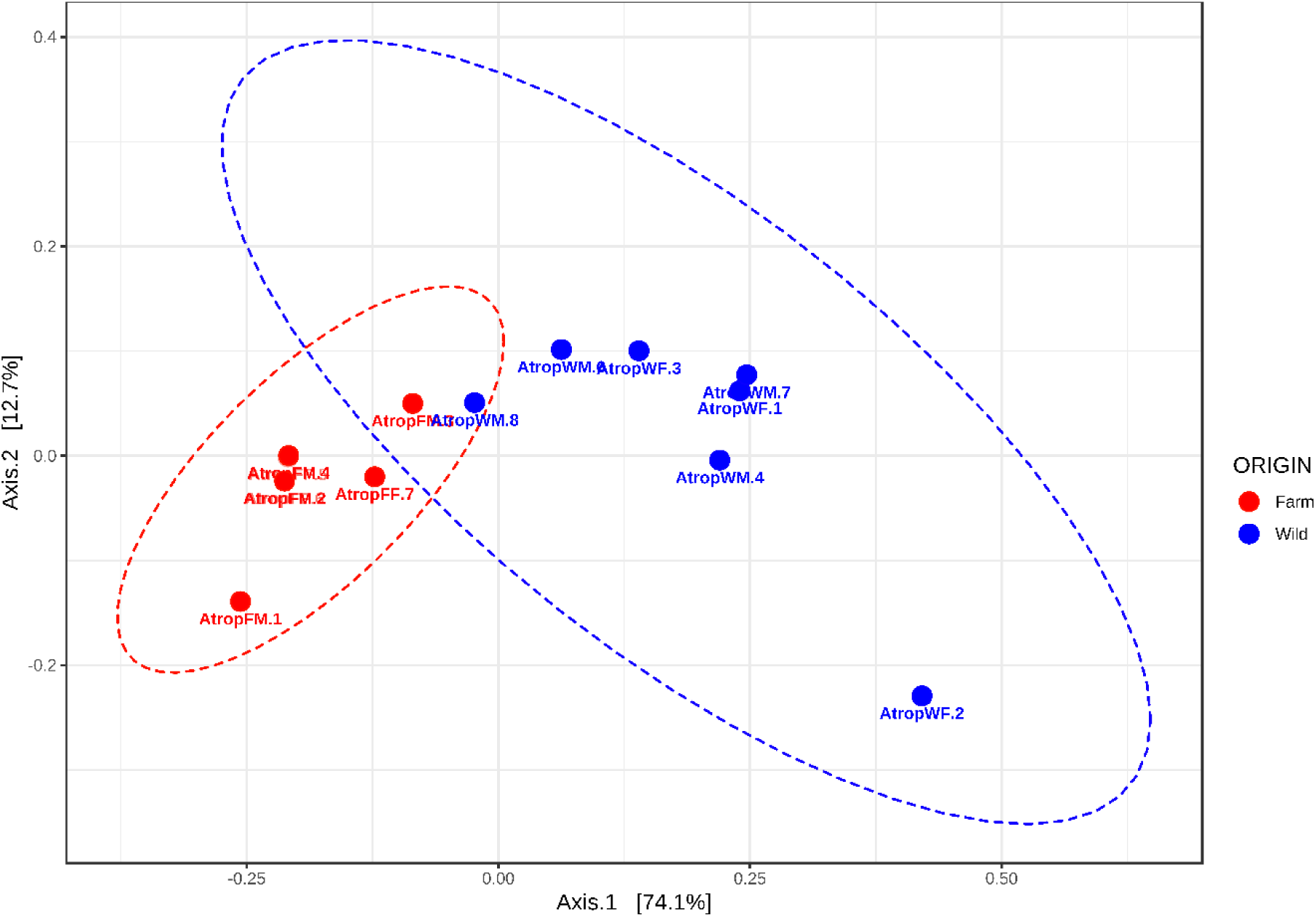
Similarity or dissimilarity coefficients of Unweighted UniFrac distances (UniFrac) by PCoA ordination analysis of the microbiota in adults’ *A. tropicus*. Blue and red circles correspond to wild and cultivated individuals, respectively

### Significant microbial components associated with the origin and sex

8 genera were identified in wild organisms and these are *Methanoculleus, Flavobacterium, Psychrobacter, Acinetobacter, Pseudomonas, Paracoccus, Massilia* and *Shewanella.* Likewise, 13 genera were identified in cultivated organisms, which are *Paludibacter, Intestinibacter, Cellulosilyticum, Odoribacter, Turicibacter, Defluviitalea, Vallitalea, Acetivibrio, Terrisporobacter, Bacteroides, Acidaminobacter, Sporacetigenium* and *Macellibacteroides.*

Specific taxa distributed differentially between wild and cultivated organisms were identified by LEfSe tool. This allows to obtain statistical differences per each taxon, where linear discriminant analysis (LDA) score barplot is shown for both conditions, cultivated and wild organisms (female & male; figure 5a), cultivated and wild organisms (female; figure 5b), male organisms, cultivated & wild (Figure 6a), cultivated female organisms (Figure 6b) and wild female organisms (Figure 6c), representing each phylum, class, order, family and genus by a histogram.

**Figure 5.**
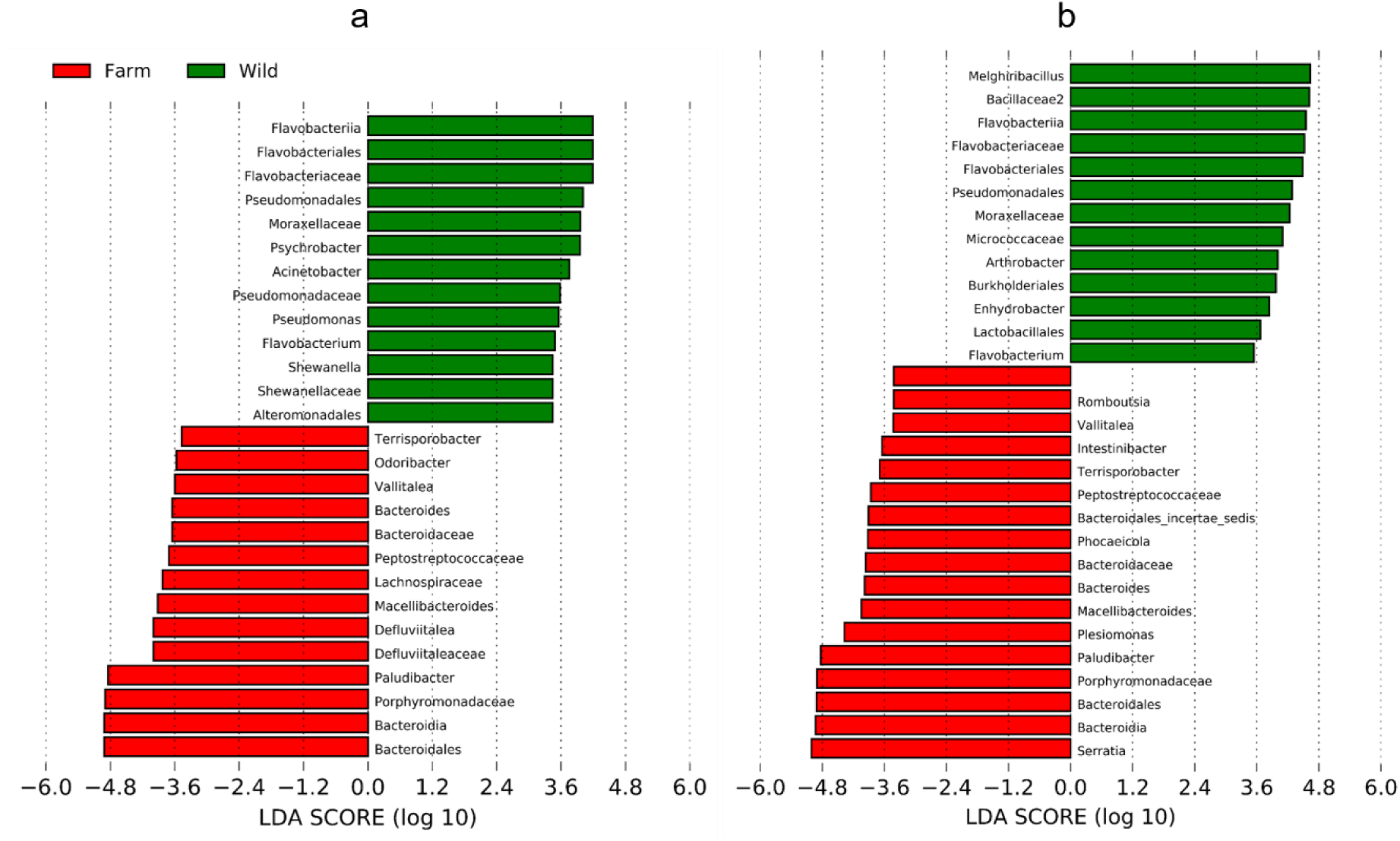
LEfSe analysis with the LDA score histogram of microbiota in adults’ *A. tropicus* composition, (a) comparison among female and male organisms, cultivated and wild, and (b) comparison among female organisms cultivated and wild. Both show statistically significant differences depending on the origin. Red and green colors indicate cultivated and wild samples, respectively.

**Figure 6.**
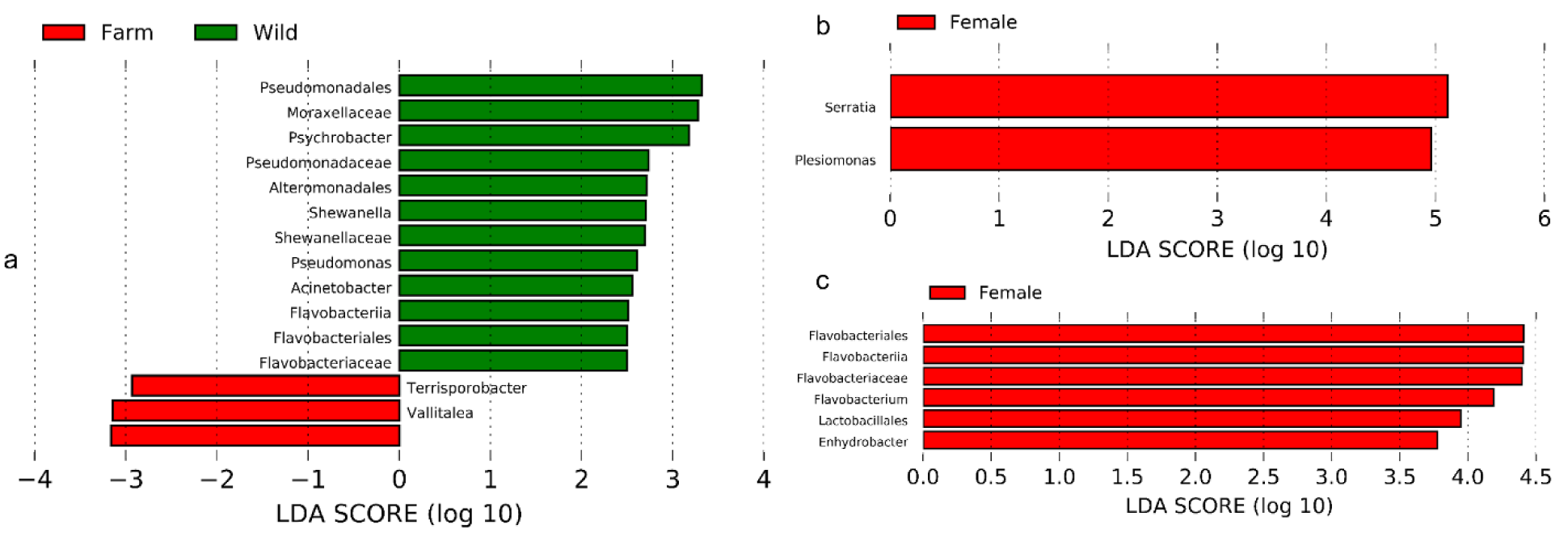
LEfSe analysis with the LDA score histogram of microbiota in adults’ *A. tropicus* composition (a) male organisms, cultivated & wild, (b) female organisms cultivated and (c) wild female. All show statistically significant differences depending on the origin.

### Phylogenetic reconstruction analysis

The sequences-base of rRNA 16S gene constructed of some organisms found in the literature with probiotic potential in fish, according to table 1, allowed us to find and identify possible organisms with probiotic potential in microbiome gut of adults’ *A. tropicus*, adjusted to 99% identity and 70% coverage. In figure 7 we obtained a phylogenetic tree based on the identified sequences of the most abundant phyla per sample, such as Fusobacteria, Proteobacteria and Firmicutes. Besides, we *Methanosarcina thermophila* and *Archaeoglobus profundus* sequences data used thermophilic methanogens and sulfate-reducing archaea, respectively, as outgroups for the root identification, considering that they may be related only distantly with identified sequences in our work.

**Figure 7.**
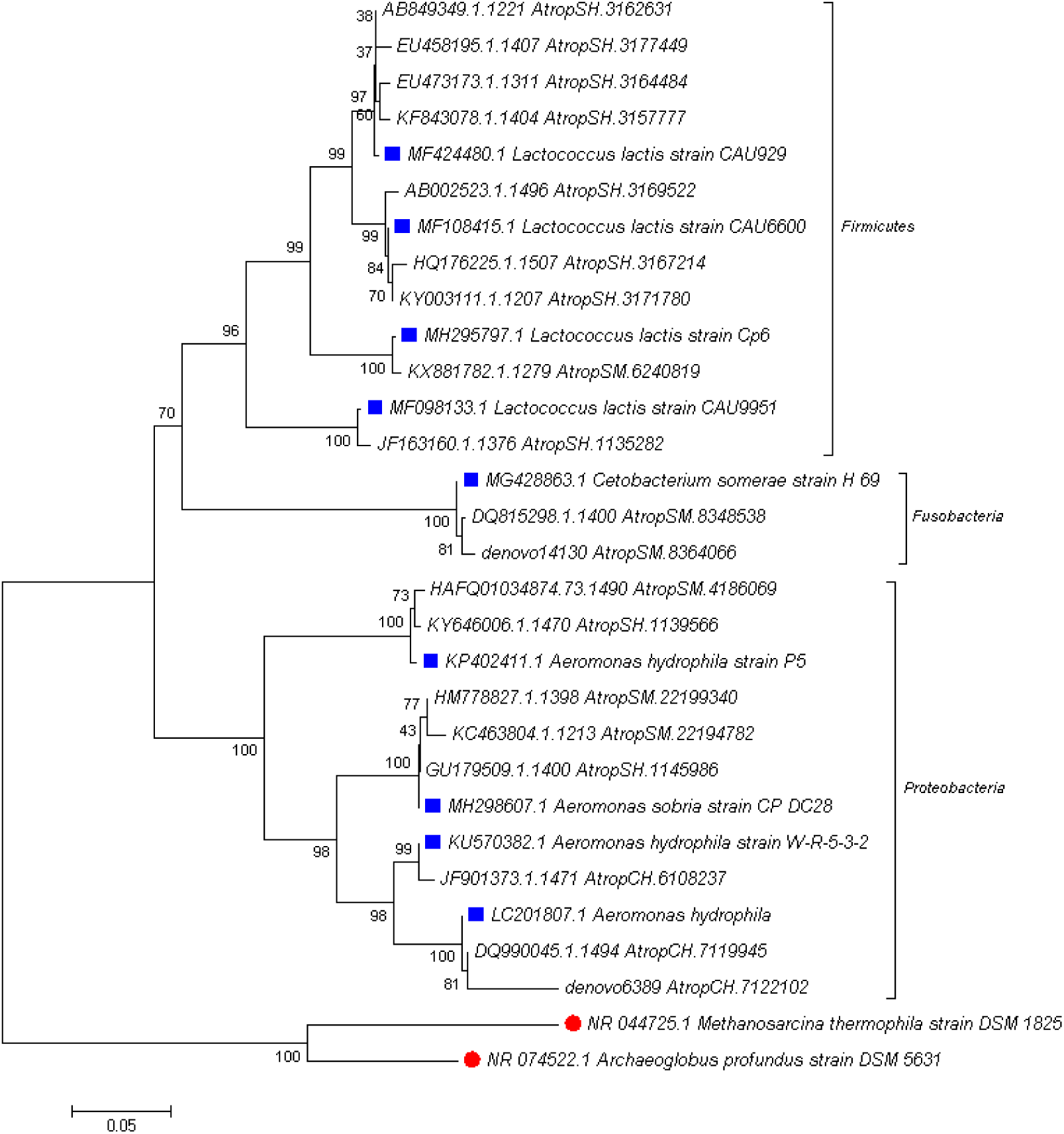
Phylogenetic tree from the rRNA 16S gene constructed of some organisms found in the literature with probiotic potential in fish (see table 1). Red circles and blue squares represent outgroup and reference sequences, respectively.

In phylogenetic tree reconstruction (Figure 7) we determined that the evolutionary conclusion of these relationships is that the two species of archaea or outgroups selected are ancestors of the nine species or ingroups identified with blue squares. Likewise, they belong to the three most dominant of phyla groups that integrate the core microbiome gut in adults’ *A. tropicus*. On the other hand, these nine species, identified by blue squares, are those that have probiotic potential.

## 4. Discussion

We obtained the 16S rRNA Gene Amplicons preparation with high-throughput sequencing using Illumina MiSeq platform by Research & Testing Labs (Lubbock, Texas, USA) services, facilitating discovery and analysis of rare taxa and detection of previously unrecognized eukaryotic and prokaryotic microbiome (Paul *et al*. 2018). In this sense, we identified and compared, by indexes attachment of multiplexing samples and readings of sequencing products for 2×300 bp paired-end, the core microbiome and microbial diversity, respectively, in the gut of adults’ *A. tropicus* by origin and sex.

Previous studies have shown that the fish gut hosts an estimated 10^7^ to 10^11^ bacteria per gram intestinal content (Nayak 2010) and the bacterial colonizers in fish gut include Proteobacteria, Fusobacteria, Firmicutes, Bacteroidetes, Actinobacteria, Clostridia, Bacilli and Verrucomicrobia (Ringø *et al*. 2006, Desai *et al*. 2012, Li *et al*. 2013a,b, Carda-Diéguez *et al*. 2014, Ingerslev *et al*. 2014a,b) with the first 4 being the most abundant, depending on environmental conditions and the host’s diet (Wang *et al*. 2017). Almost all Proteobacteria, Fusobacteria, Firmicutes, Bacteroidetes and Deinococcus-Thermus 16S rRNA sequences that we detect in CGMB of adults’ *A. tropicus* belonged to the *Cetobacterium, Edwardsiella, Serrartia* and *Clostridium sensu stricto* genres. The observed bacterial profiles in the gut of adults’ *A. tropicus*, by sex and origin, may reflect specific central microbiota, beyond the differences most likely attributable to feeding behavior. At the phylum level, almost 90 % of the total bacterial abundance only we were classified into total four phyla. Among these phyla, Proteobacteria and Fusobacteria were dominant in the fifteen fish species samples, female/male and cultivated/wild (Figs. 2, 3). The inferred physiological roles of the dominant prokaryotes are related to the metabolism of carbohydrates and nitrogenous compounds (Kormas *et al*. 2014). In contrast, dominant microbiota of marine fish is facultative anaerobes, including *Vibrio, Pseudomonas, Acinetobacter, Corynebacterium, Alteromonas, Flavobacterium* and *Micrococcus* (Onarheim *et al*. 1994, Blanch *et al*. 1997, Verner-Jeffreys *et al*. 2003).

Considering the above-mentioned, fish core gut microbiota (CGMB) can influence nutrition, growth, reproduction, general population dynamics and the host’s vulnerability to diseases thus supporting a crucial role in aquaculture practice (Ghanbari *et al*. 2015). Current DNA sequencing technologies and bioinformatic analysis have contributed towards a deeper understanding of the complex microbial communities associated to diverse habitats, including CGMB of fish in response to a variety of factors affecting the host, including temperature variations, salinity, growth stage, digestive physiology and feeding strategy (Cahill 1990, Jammal *et al*. 2017). The concept ‘core microbiota’ referred to a set of abundant microbial lineages that are shared by all individuals from the same species (Wong *et al*. 2013). The concept of CGMB has been explored both in mammalian host’s context and in freshwater fish (Turnbaugh *et al*. 2009, Roeselers *et al*. 2011, Nam *et al*. 2011, Wu *et al*. 2012, Wong *et al*. 2013).

On the other hand, Alpha diversity results indicate that the greatest gut microbiota abundance and richness is found in adults’ *A. tropicus* wildtype female, rather than adult’s wildtype males and the least gut microbiota abundance and richness is found in adults *A. tropicus*, female or male cultivated (Table 3). Ley *et al*. (2008) concluded that gut microflora of herbivorous mammals have the greatest richness and phylogenetic diversity and that both richness and phylogenetic diversity decreased among omnivores and decreased further among carnivores. We found the lowest richness and phylogenetic diversity (Table 3) in gut microbiomes from adults *A. tropicus* defined as top piscivores (carnivores; e.g., cultivated female and male *A. tropicus*). Indeed, MacFarlane *et al*. (1986) observed that farmraised fish had a simpler gut flora than their wild counterparts.

Likewise, our coefficients of similarity or dissimilarity of unweighted unifrac distances (UniFrac) by PCoA ordination analysis, indicate that gut microbiota of wild and cultivated adults *A. tropicus* are only similar by two wild samples and one cultivated and dissimilarity by most of the wild and cultivated samples (Figure 4). Principal coordinate analysis (PCoA) revealed that gut bacterial communities from adults’ *A. tropicus* by origin formed different clusters. Cultivated and wild organisms formed distinctly clusters in PCoA space (Figure 4), suggesting that the enrichment and diversity of gut microbiota are affected by the origin. This result is similar to the research of Ni *et al*. (2012) that the origin and host phylogeny they are quite related to the composition of adults’ *A. tropicus* gut bacteria. Moreover, previous studies have shown that the microbiotic diversity content in all intestinal sections depends on the fish size (Moran *et al*. 2005, Bolnick *et al*. 2014, Clements *et al*. 2014).

Numerous works have built on this, demonstrating that many species of herbivorous and omnivorous fishes contain diverse intestinal communities (Rimmer & Wiebe 1987, Clements *et al*. 1989, Clements 1991, 1997, Martínez-Díaz & Pérez-Espana 1999, Ray *et al*. 2012) and that herbivorous and detritivorous fish species harbour distinctive microbial populations (Clements *et al*. 2014). Feeding habits are also an important factor that generally influences the microbial diversity in the fish CGMB, displaying a higher diversity in the following order: carnivores> omnivores> herbivores (Ward *et al*. 2009, Larsen *et al*. 2014, Li *et al*. 2014a,b, Miyake *et al*. 2015, Liu *et al*. 2016). He *et al*. (2013) study revealed that the herbivorous carp (*Ctenopharyngodon idellus*) reported a wider variety of bacterial species than the dark carnivorous carps and Gibel (*Carassius gibelio*) which are exclusively omnivorous, and also the sea bream, under the same culture conditions. This same tendency we identify in gut core microbiome of adults *A. tropicus* by sex and origin, because although these are omnivores, they prefer carnivorous habits. (Méndez-Marín *et al*. 2012, Figure 5a). At the sex level, we observed that core microbiome is more diverse in female organisms, and particularly in wild type organisms (5b, 6a,b)

More recent sequence-based approaches show that fish hindgut microbial communities much more closely resemble those of mammals than environmental microbial communities (Fidopiastis *et al*. 2006; Sullam *et al*. 2012), especially in the prevalence of Proteobacteria, Firmicutes and Bacteroidetes (Clements *et al*. 2007, Smriga *et al*. 2010, Sullam *et al*. 2012, Ye *et al*. 2014). These findings indicate that fish, like other vertebrates, harbour specialized gastrointestinal communities (Clements *et al*. 2014). We identified Proteobacteria as the most dominant phylum in gut core microbiome of the adults’ *A. tropicus*. According to Rudi *et al*. (2018), Proteobacteria phylum is very characteristic in wildtype fish and denotes of a diet high in fat presence.

We identified Fusobacteria as the second dominant phylum in the gut of the male adults *A. tropicus*, wild and cultivated (Figure 2a). A few studies have shown Fusobacteria as dominant members of the gut microbiota of freshwater fishes (van Kessel *et al*. 2011, Di Maiuta *et al*. 2013). Fusobacteria is anaerobic, Gram-negative bacilli that produce butyrate (Bennett and Eley 1993), a short-chain fatty acid that is often the end product of the fermentation of carbohydrates including those found in mucins (Titus and Ahearn 1988, von Engelhardt *et al*. 1998). In mammals, butyrate provides many benefits to the host, including providing a majority of the energy supply to gastrointestinal cells (von Engelhardt *et al*. 1998, Collinder *et al*. 2003), enhancing mucus production, acting as an anti-carcinogen and anti-inflammatory, as well as playing a role in satiation (McBain *et al*. 1997, von Engelhardt *et al*. 1998, Andoh *et al*. 1999, Hamer *et al*. 2007). This fatty acid has been found in the gut of herbivorous and omnivorous fishes only (Clements *et al*. 1994, Clements and Choat 1995). Nuez-Ortin *et al*. (2012) demonstrated that the ability of butyric acid to inhibit potential freshwater fish pathogens, and sodium butyrate is currently sold as a food additive to promote fish health and growth. However, trials using blends of sodium butyrate and other additives have not proven beneficial (Owen *et al*. 2006, Gao *et al*. 2011).

OTUs sequences of the *Cetobacterium* genus were identified as dominant mainly in gut microbiome of the adults’ *A. tropicus*, wild and cultivated male organisms. *Cetobacterium* genus is widely identified in freshwater and warm water fish species (Tsuchiya *et al*. 2007, Larsen *et al*. 2014, Li *et al*. 2017). *Cetobacterium* genus members’ presence can perform fermentative metabolism of peptides and carbohydrates and produce vitamin B12 (cobalamin) (Larsen *et al*. 2014). *Cetobacterium* and *Bacteroides* were reported as major producers of the vitamin B12 in the intestine (Tsuchiya *et al*. 2008, Vogiatzoglou *et al*. 2009) and they were the dominant genera in grass carp’s intestine, with the abundance of more than 50 % (Li *et al*. 2015a). Animals, plants and fungi are incapable of cobalamin production and it is the only vitamin that is exclusively produced by microorganisms, particularly by anaerobes (Roth *et al*. 1996, Martens *et al*. 2002, Smith *et al*. 2007). Qi *et al*. (2017) showed that different concentrations of ammonia would affect the abundance of *Bacteroides* and *Cetobacterium* in gut fish, and the higher the concentration, the lower the abundance.

Despite we detected OTUs of Deinococcus-Thermus bacterial phyla non-dominant (0.003 ± 0.006 SD) in gut microbiome of the adults’ *A. tropicus*, wildtype males only. However, Deinococcus-Thermus species are known for their resistance to extreme stresses, such as radiation, oxidation, desiccation and high temperature (Li *et al*. 2015b). The deeply branching Deinococcus-Thermus lineage is recognized as one of the most extremophilic phylum of bacteria (Theodorakopoulos *et al*. 2013). Sequence information from Deinococcus-Thermus phylum is presently available for only a limited number of species. However, the sequenced genomes include species from both the main families (i.e., *Deinococcaceae* and *Thermaceae*) within this phylum (Griffiths & Gupta 2007). In recent years, researchers have begun using Deinococcus spp in biotechnologies and bioremediation due to their specific ability to grow and express novel engineered functions. More recently, the sequencing of several Deinococcus spp and comparative genomic analysis have provided new insight into the potential of this genus. Features such as the accumulation of genes encoding cell cleaning systems that eliminate organic and inorganic cell toxic components are widespread among Deinococcus spp. Other features such as the ability to degrade and metabolize sugars and polymeric sugars make Deinococcus spp. an attractive alternative for use in industrial biotechnology (Gerber *et al*. 2015). That is why their functional role in the gut of the adults’ *A. tropicus* deserves further research.

Bacteroidetes is a phylum composed of three large classes of Gram-negative, nonsporeforming, anaerobic or aerobic, and rod-shaped bacteria that are widely distributed in the environment, including in soil, sediments, and seawater, as well as in the guts and on the skin of animals (Ley *et al*. 2008), this was also present as the dominant phylum in adults *A. tropicus* female, wild and cultivated, usually; only this type of bacteria was detected as dominant in sample 4, cultivated male organism (Figure 2a). Likewise, a large part of the proteins synthesized by the genome of *Bacteroides*, a genus of Bacteroidetes, can break down polysaccharides and metabolize their sugars, playing a fundamental role in the degradation of complex molecules in the gut of the host. Their ability to harvest alternative energy sources from food could allow *Bacteroides* to be more competitive than other bacteria in CGMB of fish during starvation stage (Xu *et al*. 2003, Xia *et al*. 2014).

We identify *Clostridium sensu stricto* genus sequences in GMB of the adults’ *A. tropicus*, wildtype males only. This genus has also been identified in GMB of carp fish (Li *et al*. 2015a). The members of the genera *Clostridium sensu stricto* that are dominant in the intestinal microbiota of grass carp (*Ctenopharyngodon idellus*), they are also versatile in their ability to utilise various polysaccharides, such as cellulose, xylan and hemicelluloses, which constitute the major part of vegetal fibres (Uffen 1997, Uz & Ogram 2006, Li *et al*. 2015a). Others also include not only species with saccharolytic and fiber-fermenting activities but also proteolytic species (Lubbs *et al*. 2009, Pikuta *et al*. 2009, Li *et al*. 2015a).

*Serratia, Edwardsiella, Plesiomonas* and *Reyranella* are genera belonging to Enterobacteriaceae family and Proteobacteria phylum. This isolated bacterial species are facultative pathogens for fish and humans and may be isolated from fish without apparent symptoms of the disease (Walczak *et al*. 2017). *Serratia* produces serrawettin that acts as a wetting agent to reduce the surface tension of the environment (Chan *et al*. 2013). *Edwardsiella* has been isolated from tortoises (Iveson 1971), crocodiles (Iveson 1971), aquarium water (Bartlett *et al*. 1977) and from seagull roosting areas (Berg & Anderson 1972). *Edwardsiella* was isolated on several occasions during the examination of dressed catfish for *Salmonella* (Wyatt *et al*. 1979). This report provides information on the isolation, identification, and incidence of *Edwardsiella* in freshwater catfish and their environment. There are many unanswered questions regarding their importance in freshwater fish. Liu *et al*. (2015) are reports of pathogenicity of *Plesiomonas shigelloides* to fish. *P*. *shigelloides* can occur as natural intestinal flora of fish, but in case of stress conditions, the following symptoms are observed: darkening of body, hemorrhaging, fin rotting, ascitic fluid in the abdominal cavity, and lesions in internal organs (Liu *et al*. 2015). Phylogenetically, *Reyranella* genus has an evolutionary lineage within the family Rhodospirillaceae in the class Alphaproteobacteria. The type species *Reyranella massiliensis* was originally identified by Pagnier *et al*. (2011). Subsequently, Kim *et al*. (2013) emended several characteristics (e.g., nitrate reduction, respiratory quinone information) into the genus description, and more recently, Cui *et al*. 2017 found that reduction of nitrate to nitrite is variable, the predominant isoprenoid quinone is ubiquinone-10 (Q-10), major polar lipids are PME, DPG, PG, PE and one unknown aminolipid. We detect these genera mainly in gut microbiome of the adults’ *A. tropicus*, wildtype females and cultivated females and males organisms (Figures 2b, 3b). We assume that, although commonly known for their ability to cause deadly infectious diseases, this is populations of bacteria (identified as normal flora of the adults *A. tropicus*), which despite symbiotically living on and between wildtype females and cultivated females and males organisms of the adults’ *A. tropicus*, really have a positive impact on host survival (Figures 2b, 3b).

*Paludibacter* genus was found exclusively in African microbiota. This is probably due to their increased fitness to grow on polysaccharides abundant in xylan or cellulose diets (De Filippo *et al*. 2010, Thomas *et al*. 2011). We identified OTUs of *Paludibacter* genus in core gut microbiome of the adults’ *A. tropicus*, females and males cultivated organisms, only (Figure 2b).

In phylogenetic trees reconstruction (Figure 7) and according to Table 1, we identified the species *Lactococcus lactis* strains CAU929 and CAU6600, Cp6 and CAU9951, *Cetobacterium* strain H69, *Aeromonas hydrophila* strains P5 and WR-5-3-2, *Aeromonas sobria* strain CP DC28 and *Aeromonas hydrophila* with probiotic potential within the three dominant phyla in core gut microbiome of the adults *A. tropicus*.

## 5. Conclusions

We conclude that adults’ *A. tropicus* core gut microbiome is constituted by Proteobacteria, Fusobacteria, Firmicutes and Bacteroidetes phyla. Definitely, diversity and richness in core gut microbiome are higher of the female than males’ organisms and wild than cultivated organisms of the adults’ *A. tropicus*. We can also suppose that *Serratia, Edwardsiella, Plesiomonas* and *Reyranella* genera lives in symbiosis and has a positive impact on the survival of the adults’ *A. tropicus*. There is great potential in the biotechnology industry with Deinococcus-Thermus phylum, due to its extremophile capacity. Further, adults’ *A. tropicus* by sex and origin have been adapting to the environment and the diet due to their great capacity to saccharolytic, fiber fermentation and starvation activities. Finally, we can confirm that nine species can be found with the probiotic potential of 3 phyla that shape up the core gut microbiome of the adults *A. tropicus*. The CGMB of *A. tropicus* is increasingly regarded as an integral component of the host, due to important roles in the modulation of the immune system, the proliferation of the intestinal epithelium and the regulation of the dietary energy intake. Understanding the factors that influence the composition of these microbial communities is essential to health management, and the application to aquatic animals still requires basic investigation.

## Acknowledgments

The authors thanks the National Council of Science and Technology of Mexico for the thesis scholarship for postgraduate studies and the mixed scholarship granted to carry out the research stay at the University of Valencia (Spain). This research received external funding from the National Council of Science and Technology of Mexico “Strengthening of the Master’s Degree in Environmental Sciences for its Permanence in the National Register of Quality Graduates of CONACYT TAB-2014-C29-245836 key” and the project “Estudio de la fisiología digestiva en larvas y juveniles de pejelagarto (*Atractosteus tropicus*) con base en técnicas histológicas, bioquímicas y moleculares” CB-2016-01-282765 key.

## References

Al-Harbi AH, Uddin N (2004) Seasonal variation in the intestinal bacterial flora of hybrid tilapia (*Oreochromis niloticus, Oreochromis aureus*) cultured in earthen ponds in Saudi Arabia. Aquaculture 229: 37–44.

Al-Harbi AH, Uddin N (2005). Bacterial diversity of tilapia (*Oreochromis niloticus*) cultured in brackish water in Saudi Arabia. Aquaculture, 250: 566–572.

Andoh A, Bamba T, Sasaki M (1999). Physiological and anti-inflammatory roles of dietary fiber and butyrate in intestinal functions. JPEN J Parenter Enteral Nutr 23: S70–S73.

Aschfalk A, Müller W (2002) Clostridium perfringens toxin types from wild-caught Atlantic cod (Gadus morhua L.), determined by PCR and ELISA. Can J Microbiol 48:365–368.

Austin B, Austin DA (2016) Bacterial Fish Pathogens: Disease of Farmed and Wild Fish. Springer Sixth Edition. 259 pp. Doi: 10.1007/978-3-319-32674-0

Avila-Villa LA, Martínez-Porchas M, Gollas-Galván T, López-Elías JA, Mercado L, Murguia-López A, et al. (2011). Evaluation of different microalgae species and Artemia (*Artemia franciscana*) as possible vectors of necrotizing hepatopancreatitis bacteria. Aquaculture 318: 273–276.

Bartlett KH, Trust TJ, Lior H (1977). Small pet aquarium frogs as a source of Salmonella. Appl. Environ. Microbiol. 33: 1026–1029.

Bennett KW, Eley A (1993). Fusobacteria – new taxonomy and related diseases. J Med Microbiol 39: 246–254.

Berg RW, Anderson AW (1972). Salmonellae and *Edwardsiella* tarda in gull feces: a source of contamination in fish processing plants. Appl. Microbiol. 24: 501–503.

Blanch A, Alsina M, Simon M, Jofre J (1997). Determination of bacteria associated with reared turbot (*Scophthalmus maximus*) larvae. Journal of Applied Microbiology 82: 729–734.

Bolnick BI, Snowberg LK, Hirsch PE, Lauber CHL, Knight R, Caporaso JG, Svanbäck R (2014). Individuals’ diet diversity influences gut microbial diversity in two freshwater fish (threespine stickleback and Eurasian perch). Ecology Letters, 17: 979–987. https://doi.org/10.1111/ele.12301

Bondad MG, Subasinghe RP, Arthur JR (2005). Disease and health management in Asian aquaculture. Veterinary parasitology, vol. 132, no. 3-4, pp. 249–272.

Boutin S, Bernatchez L, Audet C, Derôme N (2012). Antagonistic effect of indigenous skin bacteria of brook charr (*Salvelinus fontinalis*) against Flavobacterium columnare and F. psychrophilum. Vet. Microbiol. 155: 355–361. https://doi.org/10.1016/j.vetmic.2011.09.002

Brunt J, Austin B (2005). Use of a probiotic to control lactococcosis and streptococcosis in rainbow trout, *Oncorhynchus mykiss* (Walbaum). J Fish Dis 28: 693–701.

Bussing WA (1998). Peces de las aguas continentales de Costa Rica. Ed. Univ. Costa Rica. 271 p.

Cahill M (1990). Bacterial flora of fishes: A review. Microbial Ecology, 19: 21–41. https://doi.org/10.1007/BF02015051

Carda-Diéguez M, Mira A, Fouz B (2014). Pyrosequencing survey of intestinal microbiota diversity in cultured sea bass (*Dicentrarchus labrax*) fed functional diets. FEMS Microbiology Ecology 87: 451–459.

Chan XY, Chang CY, Hong KW, Tee KK, Yin WF, Chan KG (2013). Insights of biosurfactant producing *Serratia marcescens* strain W2.3 isolated from diseased tilapia fish: a draft genome analysisGut Pathogens, 5: 29. https://doi.org/10.1186/1757-4749-5-29

Clements KD, Sutton DC, Choat JH (1989). The occurrence and characteristics of unusual protistan symbionts from surgeonfishes (F. Acanthuridae) of the Great Barrier Reef, Australia. Marine Biology, 102: 403–412.

Clements KD (1991). Endosymbiotic communities of two herbivorous labroid fishes, *Odax cyanomelas* and *O. pullus*. Marine Biology, 106: 223–229.

Clements KD, Gleeson VP, Slaytor M (1994). Shortchain fatty-acid metabolism in temperate marine herbivorous fish. J Comp Physiol B. 164: 372–377.

Clements KD Choat JH (1995). Fermentation in tropical marine herbivorous fishes. Physiol Zool. 68: 355–378.

Clements KD (1997). Fermentation and gastrointestinal microorganisms in fishes. In: Gastrointestinal Microbiology. Vol. 1: Gastrointestinal Ecosystems and Fermentations (eds Mackie RI, White BA), pp. 156–198. Chapman and Hall, New York.

Clements KD, Pasch IBY, Moran D, Turner SJ (2007). Clostridia dominate 16S rRNA gene libraries prepared from the hindgut of temperate marine herbivorous Fishes. Mar. Biol. 150: 1431–1440. https://doi.org/10.1007/s00227-006-0443-9

Clements KD, Angert ER, Montgomery WL, Choat JH (2014). Intestinal microbiota in fishes: what’s known and what’s not. Molecular Ecology, 23: 1891–1898. https://doi.org/10.1111/mec.12699

Collinder E, Bjornhag G, Cardona M, Norin E, Rehbinder C, Midtvedt T (2003). Gastrointestinal host-microbial interactions in mammals and fish: comparative studies in man, mice, rats, pigs, horses, cows, elk, reindeer, salmon and cod. Microb Ecol Health Dis 15: 66–78.

Cui Y, Chun SJ, Ko SR, Lee HG, Srivastava A, Oh HM, Ahn CY (2017). *Reyranella aquatilis* sp. nov., an alphaproteobacterium isolated from a eutrophic lake. Int J Syst Evol Microbiol, 67: 3496–350. https://doi.org/10.1099/ijsem.0.00215

De Filippo C, Cavalieri D, Di Paola M, Ramazzotti M, Poullet JB, Massart S, Collini S, Pieraccini G, Lionetti P (2010). Impact of diet in shaping gut microbiota revealed by a comparative study in children from Europe and rural Africa. Proc. Natl. Acad. Sci. U.S.A. 107: 14691–14696.

Desai AR, Links MG, Collins SA, Mansfield GS, Drew MD, Van Kessel AG et al. (2012). Effects of plant-based diets on the distal gut microbiome of rainbow trout (*Oncorhynchus mykiss*). Aquaculture 350: 134–142.

Di Maiuta N, Schwarzentruber P, Schenker M, Schoelkopf J (2013). Microbial population dynamics in the faeces of wood-eating loricariid catfishes. Lett Appl Microbiol 56, 401–407.

von Engelhardt W, Bartels J, Kirschberger S, Duttingdorf HDMZ, Busche R (1998). Role of short-chain fatty acids in the hind gut. Vet Q 20: S52–S59.

FAO (2014). The State of World Fisheries and Aquaculture. Roma.

FAO (2017). The State of World Fisheries and Aquaculture. Roma.

Ferguson RL, Buckley E, Palumbo A (1984). Response of marine bacterioplankton to differential filtration and confinement. Appl Environ Microbiol 47: 49–55.

Fidopiastis P, Bezdek D, Horn M, Kandel J (2006). Characterizing the resident, fermentative microbial consortium in the hindgut of the temperate-zone herbivorous fish, *Hermosilla azurea* (Teleostei: Kyphosidae). Mar Biol, 148: 631–642. https://doi.org/10.1007/s00227-005-0106-2

Flores-Nava A, Brown A (2010). Peces nativos de agua dulce de América del Sur de interés para la acuicultura: Una síntesis del estado de desarrollo tecnológico de su cultivo. Serie Acuicultura en Latinoamérica, FAO, 1: 3.

Frías-Quinatana CA, Marquez-Couturier G, Alvarez-González CA, Tovar-Ramírez D, Nolasco-Soria H, Galaviz-Espinosa M, Martínez-García R, Camarillo-Coop S, Martínez-Yáñez R, Gisbert E (2015). Development of digestive tract and enzyme activities during the early ontogeny of the tropical gar *Atractosteus tropicus*. Fish Physiol Biochem 41:1075–1091.

Gao YL, Storebakken T, Shearer KD, Penn M, Overland M (2011). Supplementation of fishmeal and plant protein-based diets for rainbow trout with a mixture of sodium formate and butyrate. Aquaculture, 311: 233–240.

Gerber E, Bernard R, Castang S, Chabot N, Coze F, Dreux-Zigha A, Hauser E, Hivin P, Joseph P, Lazarelli C, Letellier G, Olive J, Leonetti JP (2015). *Deinococcus* as new chassis for industrial biotechnology: biology, physiology and tools. Journal of Applied Microbiology, 119: 1–10. https://doi.org/10.1111/jam.12808

Ghanbari M, Kneifel W, Domig KJ (2015). A new view of the fish gut microbiome: Advances from next-generation sequencing. Aquaculture 448: 464–475. https://doi.org/10.1016/j.aquaculture.2015.06.033

Gianoulis TA, Raes J, Patel PV, Bjornson R, Korbel JO, Letunic I, et al. (2009). Quantifying environmental adaptation of metabolic pathways in metagenomics. Proceedings of the National Academy of Sciences, 106: 1374–1379.

Givens CE (2012). A fish tale: comparison of the gut microbiome of 15 fish species and the influence of diet and temperature on its composition. Thesis of Philosophy. University of Georgia. Pp 10–229.

Griffiths E, Gupta RS (2007). Identification of signature proteins that are distinctive of the *Deinococcus*-*Thermus* phylum. International Microbiology, 10: 201–208. https://doi.org/10.2436/20.1501.01.28

Hamer H, Jonkers D, Troost FJ, Bast A, Vanhoutvin S, Venema K, Brummer RJ (2007). Butyrate modulates oxidative stress in the colonic mucosa of healthy humans. Gastroenterology 132, A79.

Head IM, Saunders JR, Pickup RW (1998) Microbial evolution, diversity, and ecology: a decade of ribosomal RNA analysis of uncultivated microorganisms. Microb Ecol 35: 1–21.

Ingerslev HC, Strube ML, von Gersdorff Jørgensen L, Dalsgaard I, Boye M, Madsen L (2014a). Diet type dictates the gut microbiota and the immune response against Yersinia ruckeri in rainbow trout (*Oncorhynchus mykiss*). Fish & Shellfish Immunology 40: 624–633.

Ingerslev HC, von Gersdorff Jørgensen L, Strube ML, Larsen N, Dalsgaard I, Boye M et al. (2014b). The development of the gut microbiota in rainbow trout (*Oncorhynchus mykiss*) is affected by first feeding and diet type. Aquaculture 424: 24–34.

Irianto A, Austin B (2002a). Use of probiotics to control furunculosis in rainbow trout, *Oncorhynchus mykiss* (Walbaum). Journal of Fish Diseases, 25: 333–342.

Irianto A, Austin B (2002b). Use of probiotics to control furunculosis in rainbow trout, *Oncorhynchus mykiss* (Walbaum). J. Fish Dis. 25: 1–10. https://doi.org/10.1046/j.1365-2761.2002.00375.x

Iveson JB (1971). Strontium chloride B and E. E. broth media for the isolation of *Edwardsiella, Salmonella* and Arizona species from tiger snakes. J. Hyg. 69: 323–330.

Izvekova GI, Izvekov E, Plotnikov A (2007). Symbiotic microflora in fishes of different ecological groups. Biol Bull 34: 610–618.

Jammal A, Bariche M, zu Dohna H, Kambris Z (2017). Characterization of the Cultivable Gut Microflora in Wild-Caught Mediterranean Fish Species. Current Nutrition & Food Science, 13:147–154. https://doi.org/10.2174/1573401313666170216165332

Joborn A, Dorsch M, Olsson JC, Westerdahl A, Kjelleberg S (1999). *Carnobacterium inhibens* sp nov., isolated from the intestine of Atlantic salmon (Salmo salar). Int. J. Syst. Bacteriol. 49: 1891–1898.

van Kessel MA, Dutilh BE, Neveling K, Kwint MP, Veltman MP, Flik G, Jetten MS, Klaren PH et al. (2011). Pyrosequencing of 16S rRNA gene amplicons to study the microbiota in the gastrointestinal tract of carp (*Cyprinus carpio* L.). AMB Express 1, 41

Kim SJ, Ahn JH, Lee TH, Weon HY, Hong SB et al. (2013). *Reyranella soli* sp. nov., isolated from forest soil, and emended description of the genus *Reyranella Pagnier* et al. 2011. Int J Syst Evol Microbiol, 63: 3164–3167.

Klindworth A, Pruesse E, Schweer T, Peplies J, Quast Ch, Horn M, Oliver Glöckner FO (2013). Evaluation of general 16S ribosomal RNA gene PCR primers for classical and next-generation sequencing-based diversity studies. Nucleic Acids Research, 41:1–11. https://doi.org/10.1093/nar/gks808

Kormas KA, Meziti A, Mente E, Frentzos A (2014). Dietary differences are reflected on the gut prokaryotic community structure of wild and commercially reared sea bream (*Sparus aurata*). MicrobiologyOpen, 3(5): 718–728. https://doi.org/10.1002/mbo3.202

Larsen AM, Mohammed HH, Arias CR (2014). Characterization of the gut microbiota of three commercially valuable warmwater fish species. Journal of Applied Microbiology 116: 1396–1404.

Ley RE, Hamady M, Lozupone C, Turnbaugh PJ and others (2008). Evolution of mammals and their gut microbes. Science 320: 1647–1651. https://doi.org/10.1126/science.1155725

Li J, Ni J, Li X, Yan Q, Yu Y (2013a). Relationship between gastrointestinal bacterial structure and development of *Silurus soldatrovi meridodalis*. Chen. Acta Hydrobiologica Sinica 37: 613–619.

Li X, Yan Q, Xie S, Hu W, Yu Y, Hu Z (2013b). Gut microbiota contributes to the growth of fast-growing transgenic common carp (*Cyprinus carpio* L.). PLoS One 8: e64577.

Li J, Ni J, Wang C, Li X, Wu S, Zhang T et al. (2014a). Comparative study on gastrointestinal microbiota of eight fish species with different feeding habits. Journal of Applied Microbiology 117: 1750–1760.

Li X, Zhu Y, Yan Q, Ringø E, Yang D (2014b). Do the intestinal microbiotas differ between paddlefish (Polyodon spathala) and bighead carp (*Aristichthys nobilis*) reared in the same pond? Journal of Applied Microbiology 117: 1245–1252.

Li T, Long M, Gatesoupe FJ, Zhang Q, Li A, Gong X (2015a). Comparative Analysis of the Intestinal Bacterial Communities in Different Species of Carp by Pyrosequencing. Microb Ecol., Microb Ecol., 69:25–36. DOI 10.1007/s00248-014-0480-8.

Li H, Zhong Q, Wirth S, Wang W, Hao Y, Wu S, Zou H, Li W, Wang G (2015b). Diversity of autochthonous bacterial communities in the intestinal mucosa of grass carp (*Ctenopharyngodon idellus*) (Valenciennes) determined by culture-dependent and culture-independent techniques. Aquaculture Research, 46: 2344–2359. https://doi.org/10.1111/are.12391

Li T, Li H, Gatesoupe FJ, She R, Lin Q, Yan X, Li J, Li X (2017). Bacterial Signatures of “Red-Operculum” Disease in the Gut of Crucian Carp (*Carassius auratus*). Microb Ecol., 74: 510–521. htpps://doi.org/10.1007/s00248-017-0967-1

Liu Z, Ke X, Lu M, Gao F, Cao J, Zhu H, Wang M (2015). Identification and pathological observation of a pathogenic *Plesiomonas shigelloides* strain isolated from cultured tilapia (*Oreochromis niloticus*). Wei Sheng Wu Xue Bao, 55, 96–106.

Liu H, Guo X, Gooneratne R, Lai R, Zeng C, Zhan F et al. (2016). The gut microbiome and degradation enzyme activity of wild freshwater fishes influenced by their trophic levels. Scientific Reports 6: 24340.

Lubbs D, Vester B, Fastinger N, Swanson K (2009). Dietary protein concentration affects intestinal microbiota of adult cats: a study using DGGE and qPCR to evaluate differences in microbial populations in the feline gastrointestinal tract. J Anim Physiol Anim Nutr 93:113– 121

MacFarlane RD, McLaughlin JJ, Bullock GL (1986). Quantitative and qualitative studies of gut flora in striped bass from estuarine and coastal marine environments. J Wildl Dis 22: 344–348.

Márquez-Couturier G, Álvarez-González CA, Contreras-Sánchez WM, Hernández-Vidal U, Hernández-Franyutti AA, Mendoza-Alfaro RE, Aguilera-González C, García-Galano T, Civera-Cerecedo R & Goytortua-Bores E (2006). Avances en la alimentación y nutrición de pejelagarto *Atractosteus tropicus*. p. 446–523. *In:* Cruz Suárez, L. E., D. Ricque-Marie, M. Tapia-Salazar, M.G. Nieto-López, D.A. Villarreal-Cabazos, A.C. Puello & A. García-Ortega (Eds.). Avances en Nutrición Acuícola VIII. VIII Simposium Internacional de Nutrición Acuícola. 15 al 17 Noviembre. Universidad Autónoma de Nuevo León, Monterrey, Nuevo León, México.

Márquez-Couturier G, Vázquez-Navarrete CJ (2015). Estado de arte de la biología y cultivo de pejelagarto (*Atractosteus tropicus*). Agroproductividad 3:44–51. ISSN 24487546.

Márquez-Couturier G, Vázquez-Navarrete CJ, Contreras-Sánchez WM, Alvarez-González CA (2015). Acuicultura tropical sustentable: Una estrategia para la producción y conservación del pejelagarto (*Atractosteus tropicus*) en Tabasco, México. 2nd edition. Colección José Narciso Rovirosa. Villahermosa, Tabasco, Mexico. 87 p.

Martens JH, Barg H, Warren MJ, Jahn D (2002). Microbial production of vitamin B12. Appl Microbiol Biotechnol, 58: 275–285.

Martin-Antonio B, Manchado M, Infante C, Zerolo R, Labella A, Alonso C, Borrego JJ (2007) Intestinal microbiota variation in Senegalese sole (*Soleasenegalensis*) under different feeding regimes. Aquacult Res 38: 1213–1222

Martínez-Cruz P, Ibáñez AL, Monroy-Hermosillo O, Ramırez-Saad HC (2012). Use of Probiotics in Aquaculture. ISRN Microbiology, 2012:13, ID 916845. https://doi.org/10.5402/2012/916845

Martínez-Díaz SF, Pérez-Espana H (1999) Feasible mechanisms for algal digestion in the king angelfish. Journal of Fish Biology, 55: 692–703.

McBain JA, Eastman A, Nobel CS Mueller GC (1997). Apoptotic death in adenocarcinoma cell lines induced by butyrate and other histone deacetylase inhibitors. Biochem Pharmacol, 53: 1357– 1368.

Méndez-Marín O, Hernández-Franyutti AA, Álvarez-González CA, Contreras-Sánchez WM, Uribe-Aranzábal, MC (2012). Histología del ciclo reproductor de hembras del pejelagarto *Atractosteus tropicus* (Lepisosteiformes: Lepisosteidae) en Tabasco, México. Revista de Biología Tropical. Universidad de Costa Rica, Costa Rica. 60:1857–1871. ISSN: 0034-7744.

Miller RR, Minckley WL & Norris SM (2005). Freshwater fishes of Mexico. University of Chicago press, Chicago and London. 490 p.

Miyake S, Ngugi DK, Stingl U (2015). Diet strongly influences the gut microbiota of surgeonfishes. Molecular Ecology 24: 656–672.

Moran D, Turner SJ, Clements KD (2005). Ontogenetic Development of the Gastrointestinal Microbiota in the Marine Herbivorous Fish *Kyphosus sydneyanus*. Microbial Ecology, 49: 590– 597. https://doi.org/10.1007/s00248-004-0097-4

Nam YD, Jung MJ, Roh SW, Kim MS, Bae JW (2011). Comparative analysis of Korean human gut microbiota by barcoded pyrosequencing. PLoS One 6:e22109. htpps://doi.org/10.1371/journal.pone.0022109

Nayak SK (2010). Role of gastrointestinal microbiota in fish. Aquaculture Research 41: 1553–1573.

Nelson SJ (2006). Fishes of the World. 4th Edition. A Wiley-Interscience publication. EEUU. 601 p.

Newman JT Jr, Cosenza BJ, Buck JD (1972). Aerobic microflora of the bluefish (*Pomatomus saltatrix*) intestine. J Fish Res Board Can 29:333–336.

Ni J, Yu Y, Zhang T, Gao L (2012). Comparison of intestinal bacterial communities in grass carp, *Ctenopharyngodon idellus*, from two different habitats. Chinese Journal of Oceanology and Limnology, 30: 757–765. https://doi.org/10.1007/s00343-012-1287-4

Nielsen HB, Almeida M, Juncker AS, Rasmussen S, Li J, Sunagawa S, et al. (2014). Identification and assembly of genomes and genetic elements in complex metagenomic samples without using reference genomes. Nature Biotechnology 32: 822– 828.

Nuez-Ortin WG, Prado S, Toranzo AE (2012). Antimicrobial properties of butyric acid and other organic acids against pathogenic bacteria affecting the main aquatic species. In AQUA 2012. Prague.

Núñez de la Rosa MG (2011). Evaluación preliminar de las poblaciones bacterianas asociadas al tracto intestinal de la tilapia (*Oreochromis niluticus*) expuestas a aceites esenciales de orégano en la dieta. Tesis de Maestría. Facultad de Ciencias. Universidad Nacional de Colombia. Pp 1–87.

Onarheim AM, Wiik R, Burghardt J, Stackebrandt E (1994). Characterization and identification of two *Vibrio* species indigenous to the intestine of fish in cold sea water; description of *Vibrio iliopiscarius* sp. nov. Systematic and Applied Microbiology 17: 370–379.

Owen MA, Waines P, Bradley G, Davies S (2006). The effect of dietary supplementation of sodium butyrate on the growth and microflora of *Clarias gariepinus* (Burchell 1822). In Proceedings of the XII International Symposium Fish Nutrition and Feeding ed. Deneri, O.E. pp. 301. Biarritz, France

Pagnier I, Raoult D, La Scola B (2011). Isolation and characterization of *Reyranella massiliensis* gen. nov., sp. nov. from freshwater samples by using an amoeba co-culture procedure. Int J Syst Evol Microbiol, 61: 2151–2154.

Paul F, Otte J, Schmitt I, Dal Grande1 F (2018). Comparing Sanger sequencing and high-throughput metabarcoding for inferring photobiont diversity in lichens. Scientific Reports, 8: 8624. https://doi.org/10.1038/s41598-018-26947-8

Pikuta EV, Hoover RB, Marsic D, Whitman WB, Lupa B, Tang J, Krader P (2009). *Proteocatella sphenisci* gen. nov., sp. nov., a psychrotolerant, spore-forming anaerobe isolated from penguin guano. Int J Syst Evol Microbiol 59: 2302–2307.

Pirarat N, Kobayashi T, Katagiri T, Maita M, Endo M (2006). Protective effects and mechanisms of a probiotic bacterium *Lactobacillus rhamnosus* against experimental *Edwardsiella tarda* infection in tilapia (*Oreochromis niloticus*). Vet Immunol Immunopathol, 113: 339–347.

Qi XZ, Xue MY, Yang SB, Zha JW, Wang GX, Ling F (2017). Ammonia exposure alters the expression of immune-related and antioxidant enzymes-related genes and the gut microbial community of crucian carp (*Carassius auratus*). Fish & Shellfish Immunology, 70: 485–492. https://doi.org/10.1016/j.fsi.2017.09.043

Ray AK, Ghosh K, Ringø E (2012). Enzyme-producing bacteria isolated from fish gut: a review. Aquaculture Nutrition, 18: 465– 492.

Reséndez-Medina A, Salvadores-Baledón ML (1983). Contribución al conocimiento de la biología del pejelagarto *Lepisosteus tropicus* (Gill) y la tenguayaca *Petenia splendida* Günther, del estado de Tabasco. Biótica 8: 413–426.

Rimmer DW, Wiebe RJ (1987). Fermentative microbial digestion in herbivorous fishes. Journal of Fish Biology, 31:229–236.

Ringø E, Strøm E, Tabachek JA (1995). Intestinal microflora of salmonids: a review. Aquacult Res. 26: 773–789.

Ringø E, Sperstad S, Myklebust R, Refstie S, Krogdahl Å (2006). Characterisation of the microbiota associated with intestine of Atlantic cod (*Gadus morhua* L.): the effect of fish meal, standard soybean meal and a bioprocessed soybean meal. Aquaculture 261: 829–841.

Roeselers G, Mittge EK, Stephens WZ, Parichy DM, Cavanaugh CM, Guillemin K, Rawls JF (2011). Evidence for a core gut microbiota in the zebrafish. ISME J 5: 1595–1608

Roth JR, Lawrence JG, Bobik TA (1996). Cobalamin (coenzyme B12): synthesis and biological significance. Annu Rev Microbiol, 50: 137–181.

Rudi K, Angell IL, Pope PB, Vik JO, Sandve SR, Snipen LG (2018). Stable Core Gut Microbiota across the Freshwater-to-Saltwater Transition for Farmed Atlantic Salmon. Applied and Environmental Microbiology, 84: e01974–17. https://doi.org/10.1128/AEM.01974-17

Salinas I, Cuesta A, Esteban MA, Meseguer J (2005). Dietary administration of *Lactobacillus delbriieckii* and *Bacillus subtilis*, single or combined, on gilthead sea bream cellular innate immune responses. Fish Shellfish Immunol 19: 67–77.

Skrodenyté Arbaĉiauskiené V, (2007). Enzymatic activity of intestinal bacteria in roach Rutilus rutilus L. Fish Sci, 73: 964–966.

Smriga S, Sandin SA, Azam F (2010). Abundance, diversity, and activity of microbial assemblages associated with coral reef fish guts and feces. FEMS Microbiology Ecology, 73: 31–42. https://doi.org/10.1111/j.1574-6941.2010.00879.x

Smith AG, Croft MT, Moulin M, Webb ME (2007). Plants need their vitamins too. Curr Opin Plant Biol, 10: 266–275.

Spanggaard B, Huber I, Nielsen J, Nielsen T, Appel KF, Gram L (2000). The microflora of rainbow trout intestine: a comparison of traditional and molecular identification. Aquaculture 182: 1–15.

Subasinghe RP (2005). Epidemiological approach to aquatic animal health management: opportunities and challenges for developing countries to increase aquatic production through aquaculture. Prev. Vet. Med. 67: 117–124.

Sullam KE, Essinger SD, Lozupone CA, O’Connor MP, Rosen GL, Knight R, Kilham SS, Russell JA (2012). Environmental and ecological factors that shape the gut bacterial communities of fish: a meta-analysis. Molecular Ecology, 21: 3363–3378. https://doi:10.1111/j.1365-294X.2012.05552.x

Suttle CA (2007). Marine viruses—major players in the global ecosystem. Nature Reviews Microbiology, 5: 801–812.

Tamaki H, Wright CL, Li X, Lin Q, Hwang C, Wang S, Thimmapuram J, Kamagata Y, Liu WT (2011). Analysis of 16S rRNA Amplicon Sequencing Options on the Roche/454 Next-Generation Titanium Sequencing Platform. PLOS ONE 6: e25263. https://doi.org/10.1371/journal.pone.0025263

Theodorakopoulos N, Bachar D, Christen R, Alain K, Chapon V (2013). Exploration of Deinococcus-Thermus molecular diversity by novel group-specific PCR primers. Microbiologyopen, 2: 862–72. https://doi.org/10.1002/mbo3.119

Thomas F, Hehemann JH, Rebuffet E, Czjzek M, Michel G (2011). Environmental and gut Bacteroidetes: the food connection. Front Microbiol, 2: 93. https://doi.org/10.3389/fmicb.2011.00093

Titus E, Ahearn GA (1988). Short-chain fatty-acid transport in the intestine of a herbivorous teleost. J Exp Biol 135, 77–94.

Tsuchiya C, Sakata T, Sugita H (2007). Novel ecological niche of *Cetobacterium somerae*, an anaerobic bacterium in the intestinal tracts of freshwater fish. Letters in Applied Microbiology ISSN 0266-8254.

Tsuchiya C, Sakata T, Sugita H (2008). Novel ecological niche of *Cetobacterium somerae*, an anaerobic bacterium in the intestinal tracts of freshwater fish. Letters in Applied Microbiology 46: 43– 48. https://doi.org/10.1111/j.1472-765X.2007.02258.x.

Turnbaugh PJ, Hamady M, Yatsunenko T, Cantarel BL, Duncan A, Ley RE, Sogin ML, Jones WJ, Roe BA, Affourtit JP (2009). A core gut microbiome in obese and lean twins. Nature 457: 480–484

Uffen R (1997). Xylan degradation: a glimpse at microbial diversity. J Ind Microbiol Biotechnol 19:1–6.

Uz I, Ogram AV (2006). Cellulolytic and fermentative guilds in eutrophic soils of the Florida Everglades. FEMS Microbiol Ecol 57:396–408

Verner-Jeffreys DW, Shields RJ, Bricknell IR, Birkbeck TH (2003). Changes in the gut-associated microflora during the development of Atlantic halibut (*Hippoglossus hippoglossus* L.) larvae in three British hatcheries. Aquaculture 219: 21–42.

Verschuere L, Rombaut G, Sorgeloos P, Verstraete W (2000). Probiotic Bacteria as Biological Control Agents in Aquaculture Microbiology and Molecular Biology Reviews, 64: 655–671.

Villamil L, Tafalla C, Figueras A, Novoa B (2002). Evaluation of immunomodulatory effects of lactic acid bacteria in turbot (*Scophthalmus maximus*). Clin Diagn Lab Immunol 9:1318–1323.

Vogiatzoglou A, Smith AD, Nurk E, Berstad P, Drevon CA, Ueland PM, Vollset SE, Tell GS, Refsum H (2009). Dietary sources of vitamin B-12 and their association with plasma vitamin B-12 concentrations in the general population: the Hordaland Homocysteine Study. The American Journal of Clinical Nutrition, 89: 1078–1087. https://doi.org/10.3945/ajcn.2008.26598

Wang AR, Ran C, Ringø E, Zhou ZG (2017). Progress in fish gastrointestinal microbiota research. Aquaculture 0:1–15. https://doi.org/10.1111/raq.12191

Walczak N, Puk K, Guz L (2017). Bacterial flora associated with diseased freshwater ornamental fish. J Vet Res, 61: 445–449. https://doi.org/10.1515/jvetres-2017-0070

Ward NL, Steven B, Penn K, Methe BA, Detrich WH (2009). Characterization of the intestinal microbiota of two Antarctic notothenioid fish species. Extremophiles 13: 679–685.

Weisburg WG, Barns SM, Pelletier DA Lane DJ (1991). 16S ribosomal DNA amplification for phylogenetic study. J. Bacteriol. 173: 697–703.

Wiley O (1976). The phylogeny and biogeography of fossil and recent gars (Actinopterygii: Lepisosteidae). Univ. KS. Hist. Nat. Mus. Publ. 64: 1–11.

Wong S, Waldrop T, Summerfelt S, Davidson J, Barrows F, Kenney PB, Welch T, Wiens GD, Snekvik K, Rawls JF (2013). Aquacultured rainbow trout (Oncorhynchus mykiss) possess a large core intestinal microbiota that is resistant to variation in diet and rearing density. Appl Environ Microbiol 79:4974–4984

Wu S, Wang G, Angert ER, Wang W, Li W, Zou H (2012). Composition, diversity, and origin of the bacterial community in grass carp intestine. PLoS One 7:e30440. doi: 10.1371/journal.pone.0030440

Wyatt LE, Nickelson R, Vanderzant C (1979). Occurrence and control of Salmonella in freshwater catfish. J. Food Sci. 44: 1067–1069.

Xia JH, Lin G, Fu GH, Wan ZY, Lee M, Wang L, Liu XJ, Yue GH (2014). The intestinal microbiome of fish under starvation. BMC Genomics, 15:266. http://doi.org/10.1186/1471-2164-15-266

Xu J, Bjursell MK, Himrod J, Deng S, Carmichael LK, Chiang HC, Hooper LV, Gordon JI (2003). A genomic view of the human-Bacteroides thetaiotaomicron symbiosis. Science, 299:2074– 2076.

Ye L, Amberg J, Chapman D, Gaikowski M, Liu WT (2014). Fish gut microbiota analysis differentiates physiology and behaviour of invasive Asian carp and indigenous American fish. ISME Journal, 8: 541–551. https://doi.org/10.1038/ismej.2013.181

