## Supplementary material for "Gut Microbiome Analysis In Adult Tropical Gars (*Atractosteus tropicus*)": TABLES

**Table 1.** Bibliographic information obtained from certain genera with probiotic potential in fish

| Bacteria | Function | Description | Fish | Reference |
|---|---|---|---|---|
| *Aeromonas hydrophila* | probiotic | antagonist | *Oncorhynchus mykiss* | Irianto A, Austin B (2002b). Use of probiotics to control furunculosis in rainbow trout, *Oncorhynchus mykiss* (Walbaum). J. Fish Dis. 25:1–10. doi: 10.1046/j.1365-2761.2002.00375.x |
| *Aeromonas salmonicida* | pathogen | etiological agent of septicemic disease | *Oncorhynchus mykiss* | |
| *Rhodococcus qingshengii* | probiotic | antagonist | *Salevinus fontinalis* | Boutin S, Bernatchez L, Audet C, Derôme N (2012). Antagonistic effect of indigenous skin bacteria of brook charr (*Salvelinus fontinalis*) against Flavobacterium columnare and F.psychrophilum. Vet. Microbiol. 155: 355–361. doi: 10.1016/j.vetmic.2011.09.002 |
| *Flavobacterium psychrophilum* | pathogen | systemic infectious agent | *Salevinus fontinalis* | |
| *Cetobacterium somerae* | probiotic | vitamin B12 production | various | Tsuchiya C, Sakata T, Sugita H (2008). Novel ecological niche of *Cetobacterium somerae*, an anaerobic bacterium in the intestinal tracts of (1), 43–48.reshwater fish. Lett. Appl. Microbiol. 46:43–48. doi:10.1111/j.1472-765X.2007.02258.x |
| *Vibrio fluvialis* | probiotic | Immune stimulation and improved survival after challenge with *Aeromonas salmonicida* | *Oncorhynchus mykiss* | Irianto A & Austin B (2002a). Use of probiotics to control furunculosis in rainbow trout *Oncorhynchus mykiss* (Walbaum). J Fish Dis 25: 333–342 |
| *Lactococcus lactis CECT 539* | probiotic | Immune stimulation | *Scophthalmus maximus* | Villamil L, Tafalla C, Figueras A, Novoa B (2002). Evaluation of immunomodulatory effects of lactic acid bacteria in turbot (*Scophthalmus maximus*). Clin Diagn Lab Immunol 9:1318–1323. |
| *Lactobacillus delbrueckii CECT 287* | probiotic | Immune stimulation | *Sparus aurata* | Salinas I, Cuesta A, Esteban MA, Meseguer J (2005). Dietary administration of *Lactobacillus delbriieckii* and *Bacillus subtilis*, single or combined, on gilthead sea bream cellular innate immune responses. Fish Shellfish Immunol 19: 67–77. |
| *Bacillus subtilis CECT 35* | probiotic | Immune stimulation | *Sparus aurata* | |
| *Aeromonas sobria GC2* | probiotic | Immune stimulation and improved survival after challenge with *Lactococcus garvieae* and *Streptococcus iniae* | *Oncorhynchus mykiss* | Brunt J, Austin B (2005). Use of a probiotic to control lactococcosis and streptococcosis in rainbow trout, *Oncorhynchus mykiss* (Walbaum). J Fish Dis 28: 693–701. |
| *Lactobacillus rhamnosus ATCC 53103* | probiotic | Immune stimulation and improved survival after challenge with *Edwardsiella tarda* | *Oreochromis niloticus* | Pirarat N, Kobayashi T, Katagiri T, Maita M, Endo M (2006). Protective effects and mechanisms of a probiotic bacterium *Lactobacillus rhamnosus* against experimental *Edwardsiella tarda* infection in tilapia (*Oreochromis niloticus*). Vet Immunol Immunopathol, 113: 339–347. |
| *Carnobacterium inhibens* | probiotic | antibacterial activity against fish pathogens | *Salmo salar* | Joborn A, Dorsch M, Olsson JC, Westerdahl A, Kjelleberg S (1999). *Carnobacterium inhibens* sp nov., isolated from the intestine of Atlantic salmon (Salmo salar). Int. J. Syst. Bacteriol. 49:1891–1898. |
| *Enterobacter amnigenus* | probiotic | increased resistance toward *Flavobacterium psychrophilum* | *Oncorhynchus mykiss* | Irianto A, Austin B (2002b). Use of probiotics to control furunculosis in rainbow trout, *Oncorhynchus mykiss* (Walbaum). J. Fish Dis. 25:1–10. doi: 10.1046/j.1365-2761.2002.00375.x |

**Table 2**. Total reads by sex/origin and wild/cultivated

| Organisms | Reads | |
|---|---|---|
| | Wild | Cultivated |
| Female | 99,513 | 59,539 |
| Male | 137,792 | 67,891 |
| Subtotal | 237,305 | 127,430 |
| Total | 364,735 | |

**Table 3**. Alpha diversity analysis per sample, sex/origin and origin

| Sample | Chao1 | S.D. (Chao1) | Obs. OTUs | S.D. (Obs. OTUs) | P.D. | S.D. (P.D.) | Shannon | S.D. (Shannon) |
|---|---|---|---|---|---|---|---|---|
| AtropFF.5 | 250.20 $^{h}$ | 57.17 | 179 $^{fg}$ | 53.77 | 6.09 $^{fg}$ | 1.27 | 4.12 $^{k}$ | 0.12 |
| AtropFF.6 | 229.41 $^{g}$ | 56.92 | 154 $^{g}$ | 49.37 | 6.36 $^{g}$ | 1.52 | 3.54 $^{j}$ | 0.11 |
| AtropFF.7 | 241.60 $^{h}$ | 46.18 | 185 $^{j}$ | 51.61 | 9.16 $^{j}$ | 1.64 | 4.46 $^{m}$ | 0.13 |
| AtropFM.1 | 75.65 $^{b}$ | 22.22 | 54 $^{a}$ | 15.88 | 2.60 $^{a}$ | 0.61 | 1.50 $^{c}$ | 0.06 |
| AtropFM.2 | 144.24 $^{e}$ | 40.38 | 97 $^{d}$ | 31.52 | 4.10 $^{d}$ | 0.94 | 2.03 $^{e}$ | 0.10 |
| AtropFM.3 | 101.88 $^{d}$ | 29.58 | 69 $^{a}$ | 21.85 | 2.54 $^{a}$ | 0.51 | 2.52 $^{f}$ | 0.07 |
| AtropFM.4 | 210.05 $^{f}$ | 50.58 | 146 $^{f}$ | 46.20 | 5.94 $^{f}$ | 1.38 | 3.24 $^{h}$ | 0.11 |
| AtropWF.1 | 92.24 $^{c}$ | 29.32 | 60 $^{e}$ | 20.20 | 4.79 $^{e}$ | 1.20 | 1.18 $^{b}$ | 0.06 |
| AtropWF.2 | 456.85 $^{j}$ | 110.76 | 296 $^{k}$ | 98.54 | 13.52 $^{k}$ | 3.52 | 5.43 $^{n}$ | 0.17 |
| AtropWF.3 | 262.49 $^{i}$ | 61.28 | 186 $^{i}$ | 56.11 | 7.64 $^{i}$ | 1.67 | 4.43 $^{l}$ | 0.13 |
| AtropWM.4 | 49.84 $^{a}$ | 14.01 | 37 $^{e}$ | 11.28 | 4.60 $^{e}$ | 0.72 | 0.86 $^{a}$ | 0.06 |
| AtropWM.6 | 216.94 $^{f}$ | 69.18 | 119 $^{h}$ | 42.08 | 7.05 $^{h}$ | 3.10 | 3.29 $^{i}$ | 0.09 |
| AtropWM.7 | 141.11 $^{e}$ | 36.74 | 97 $^{c}$ | 30.10 | 3.63 $^{c}$ | 0.90 | 3.16 $^{g}$ | 0.08 |
| AtropWM.8 | 94.20 $^{cd}$ | 34.04 | 59 $^{b}$ | 18.75 | 3.31 $^{b}$ | 1.52 | 1.57 $^{d}$ | 0.07 |

Kruskal-Wallis = p<0.01

| Sex/Origin | Chao1 | S.D. (Chao1) | Obs. OTUs | S.D. (Obs. OTUs) | P.D. | S.D. (P.D.) | Shannon | S.D. (Shannon) |
|---|---|---|---|---|---|---|---|---|
| FF | 465.74 $^{a}$ | 67.21 | 402 $^{a}$ | 91.40 | 14.62 $^{a}$ | 2.47 | 4.22 $^{a}$ | 0.06 |
| FM | 374.76 $^{b}$ | 66.59 | 302 $^{b}$ | 79.96 | 9.57 $^{b}$ | 2.04 | 3.24 $^{b}$ | 0.05 |
| WF | 803.73 $^{c}$ | 120.84 | 675 $^{c}$ | 169.82 | 23.74 $^{c}$ | 4.77 | 5.57 $^{c}$ | 0.09 |
| WM | 447.04 $^{d}$ | 88.21 | 339 $^{d}$ | 98.71 | 15.49 $^{d}$ | 4.50 | 4.05 $^{d}$ | 0.05 |

Kruskal-Wallis = p<0.01

| Origin | Chao1 | S.D. (Chao1) | Obs. OTUs | S.D. (Obs. OTUs) | P. D. | S.D. (P.D.) | Shannon | S.D. (Shannon) |
|---|---|---|---|---|---|---|---|---|
| Farm | 543.24 $^{a}$ | 76.85 | 486 $^{a}$ | 105.77 | 16.29 $^{a}$ | 2.79 | 3.88 $^{a}$ | 0.07 |
| Wild | 1090.53 $^{b}$ | 178.13 | 912 $^{b}$ | 239.31 | 31.67 $^{b}$ | 7.35 | 5.35 $^{b}$ | 0.10 |

W of Mann-Whitney (Wilcoxon) p<0.01

Multiple Range Test for Kruskal-Wallis (Bonferroni post-hoc):
- Different letters = p <0.01
- Equal letters = p> 0.05

FF farmed o cultivated female
FM farmed o cultivated male
WF wildtype female
WM wildtype male
